# Dynamic regulation of the oxidative stress response by the E3 ligase TRIP12

**DOI:** 10.1101/2024.11.25.625235

**Authors:** Andrew J. Ingersoll, Devlon M. McCloud, Jenny Y. Hu, Michael Rape

## Abstract

The oxidative stress response is centered on the transcription factor NRF2 and protects cells from reactive oxygen species (ROS). While ROS inhibit the E3 ligase CUL3^KEAP1^ to stabilize NRF2 and elicit antioxidant gene expression, cells recovering from stress must rapidly reactivate CUL3^KEAP1^ to prevent reductive stress and oxeiptosis-dependent cell death. How cells restore efficient NRF2-degradation upon ROS clearance remains poorly understood. Here, we identify TRIP12, an E3 ligase dysregulated in Clark-Baraitser Syndrome and Parkinson’s Disease, as a component of the oxidative stress response. TRIP12 is a ubiquitin chain elongation factor that cooperates with CUL3^KEAP1^ to ensure robust NRF2 degradation. In this manner, TRIP12 accelerates stress response silencing as ROS are being cleared, but limits NRF2 activation during stress. The need for dynamic control of NRF2-degradation therefore comes at the cost of diminished stress signaling, suggesting that TRIP12 inhibition could be used to treat degenerative pathologies characterized by ROS accumulation.

## Introduction

Reactive oxygen species (ROS), such as superoxide anions, hydroxyl radicals, or hydrogen peroxide, disrupt cell and tissue integrity and thereby accelerate aging and neurodegeneration ^1–3^. Such pathological ROS can result from an exposure to toxins or heavy metals, but also accumulate during infection or due to mutations in mitochondrial proteins or ROS-detoxifying enzymes ^1,2^. The cytotoxic effects of ROS are exploited in the clinic during radiation and chemotherapy to eliminate tumor cells and provide cancer patients with therapeutic benefit ^4^.

To counteract deleterious ROS, cells rely on the oxidative stress response that is centered on the transcription factor NRF2 and its E3 ligase CUL3^KEAP1 5–7^. In unstressed cells, CUL3^KEAP1^ constantly ubiquitylates NRF2 to induce its proteasomal degradation and keep NRF2 levels low ^8,9^. As ROS accumulate, oxidation of Cys residues in the substrate adaptor KEAP1 prevents NRF2 turnover, which allows the transcription factor to enter the nucleus and drive antioxidant gene expression ^5,10–13^. While rapid NRF2 stabilization protects cells from ROS, efficient reactivation of CUL3^KEAP1^ during recovery is also important: a failure to restore NRF2 turnover upon ROS clearance triggers reductive stress, which impairs differentiation and causes a range of developmental and metabolic diseases ^14–19^. Persistent inhibition of KEAP1 also induces a cell death pathway referred to as oxeiptosis ^20^. In line with these observations, *KEAP1* deletion in mice causes perinatal lethality ^21^, and mutations in the *KEAP1* locus are major drivers of lung adenocarcinoma and other tumors ^22,23^. Cells must therefore balance their need to rapidly stabilize NRF2 during stress with an ability to reactivate CUL3^KEAP1^ once conditions improve. How cells establish this dynamic control of oxidative stress signaling is not fully understood.

CUL3^KEAP1^ differs from other ubiquitylation enzymes that elicit proteasomal degradation. While most E3 ligases bind their substrates very briefly ^24^, KEAP1 can stably interact with NRF2 and sequester it in the cytoplasm ^25,26^. Although ROS prevent CUL3^KEAP1^ from ubiquitylating NRF2, they do not impede the ability of KEAP1 to bind the transcription factor ^10,27,28^, and KEAP1 variants known as ‘anchor mutants’ even show increased affinity for NRF2 without triggering degradation ^29^. Moreover, hetero-bifunctional small molecules that recruit CUL3^KEAP1^ to pathological proteins usually do not elicit turnover ^30^. These results mirror other CUL3 E3 ligases that rely on distinct substrate adaptors but use the same catalytic module to monoubiquitylate stable proteins ^31–34^. These findings indicated that CUL3^KEAP1^ may not be sufficient to induce robust NRF2 degradation, but whether other ubiquitylation enzymes are required is not known.

Aside from NRF2, CUL3^KEAP1^ targets the PALB2 protein that recruits BRCA2 to sites of DNA damage ^35^. Ubiquitylation of PALB2 does not change its abundance, but prevents binding to BRCA2 to avoid homologous recombination repair in the G1 phase of the cell cycle. KEAP1 also associates with the phosphatase PGAM5 ^36^, a robust interaction that is regulated by ROS during oxeiptosis ^20^. In addition, CUL3^KEAP1^ engages p62 (SQSTM1), which controls early steps in autophagy ^37,38^. Rather than inducing its degradation, CUL3^KEAP1^ ubiquitylates p62 to release an auto-inhibitory conformation for recognition of autophagic cargo ^39^. p62 engages KEAP1 so stably that CUL3^KEAP1^ is sequestered away from other substrates ^40–42^, which impedes NRF2 turnover and elicits further p62 expression ^37,43^. Together, these observations underscored that recognition by CUL3^KEAP1^ often does not result in proteasomal turnover. How cells ensure efficient NRF2 degradation before and after oxidative stress therefore requires further investigation.

Here, we show that the E3 ligase TRIP12, which is mutated in Clark-Baraitser Syndrome and overexpressed in Parkinson’s Disease ^44–47^, is a crucial component of the oxidative stress response. TRIP12 is a ubiquitin chain elongation factor that cooperates with CUL3^KEAP1^ to decorate NRF2 with K29-linked conjugates known to drive proteasomal degradation. Chain extension by TRIP12 accelerates stress response silencing as CUL3^KEAP1^ is being restored, but restricts NRF2 function during stress. Efficient degradation of NRF2 therefore requires an additional factor, TRIP12, whose inhibition could be exploited to prolong antioxidant signaling in degenerative pathologies driven by ROS accumulation.

## Results

### CCNF is required for oxidative stress signaling in myoblasts

To understand how cells establish the dynamic nature of oxidative stress signaling, we searched for proteins that regulate NRF2 stability in differentiating myoblasts, a cell type that is dependent on precise ROS management. As seen before ^14,15,34^, KEAP1 depletion interfered with myotube formation, while reducing NRF2 had the opposite effect and improved differentiation (**Figure S1A**). Because NRF2 is required for the effects of low KEAP1 ^15^, reducing the abundance of NRF2 allowed myogenesis to occur even if cells expressed little KEAP1 (**Figure S1A**). Depleting proteins that modulate the stability or function of NRF2 might therefore restore myotube formation in the presence of low levels of KEAP1.

We had previously used this screening paradigm to probe substrate adaptors of CUL2 and CUL3 E3 ligases, which allowed us to discover the CUL2^FEM1B^-dependent reductive stress response ^15^. We now interrogated the remaining Cullin-RING E3 ligases (CUL1, CUL4A/B, CUL5) and asked if depletion of their substrate adaptors enabled myoblast differentiation in the absence of KEAP1. Strikingly, targeting the CUL1/SCF-adaptor CCNF completely restored myotube formation in cells depleted of KEAP1 (**Figure 1A**), even though reducing CCNF by itself slightly impaired this cell fate decision (**Figure S1B**). We validated these results with independent siRNAs, which confirmed that limiting the abundance of CCNF enabled the differentiation of KEAP1-depleted myoblasts (**Figure 1B**).

**Figure 1:**
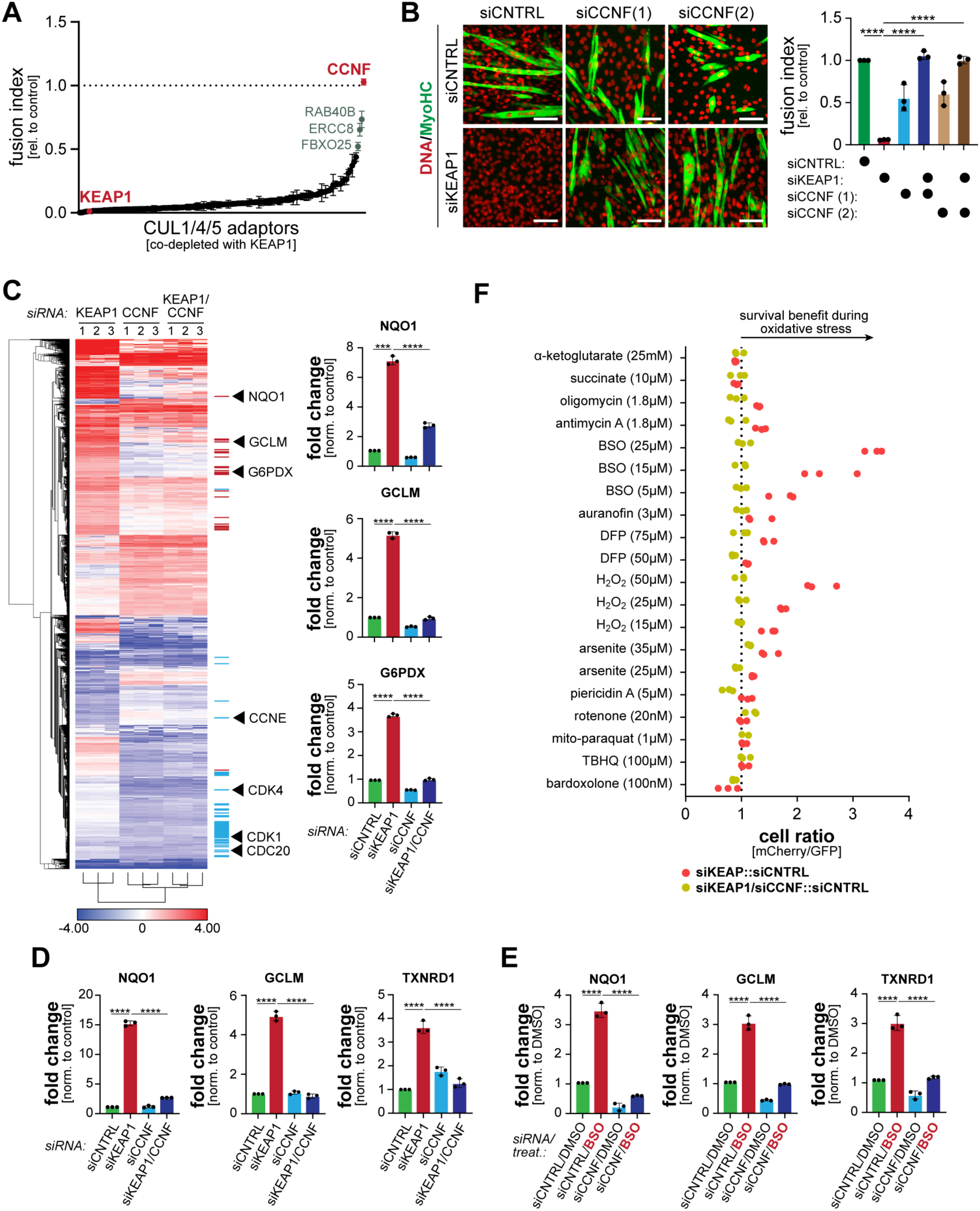
CCNF is required for oxidative stress signaling in myoblasts. **A.** A focused genetic screen identifies CCNF as a candidate regulator of oxidative stress signaling in myoblasts. C2C12 myoblasts were depleted of substrate adaptors or CUL1, CUL4, and CUL5. The same cells were depleted of KEAP1, a treatment that normally prevents myogenesis ^15^. Myoblasts were induced to differentiate, and the success of myotube formation was followed by immunostaining against MyoHC. n=2 independent screens. **B.** Validation of screen results. C2C12 myoblasts were depleted of either KEAP1, CCNF (using independent siRNAs), or both, induced to differentiate, and analyzed by immunofluorescence microscopy against MyoHC. *Left*: microscopy pictures; scale bar 100μm. *Right*: quantification of three independent experiments. **C.** RNA-sequencing analysis of C2C12 myoblasts that were depleted of either KEAP1, CCNF, or both. *Red lines:* NRF2 target genes; *blue lines:* E2F target genes. *Right*: quantification of gene expression for select NRF2 targets. **D.** C2C12 myoblasts were depleted of KEAP1, CCNF, or both, and expression of select NRF2 targets was analyzed by qPCR. **E.** C2C12 myoblasts were treated with 25μM BSO, depleted of CCNF, or both, and expression of select NRF2 targets was analyzed by qPCR. **F.** GFP-labeled control cells were mixed at a 1:1 ratio with mCherry-labeled cells depleted of KEAP1 (red dots). As indicated, CCNF was also depleted (yellow dots). Cells were exposed to increasing concentrations of oxidative stressors. After two days, the ratio of mCherry- to GFP-labeled cells was determined by flow cytometry. n=3 independent experiments. See also Figure S1.

In the absence of KEAP1, myoblasts activate NRF2 and thereby induce antioxidant proteins that interfere with differentiation ^15^. To test if CCNF acted through NRF2, we subjected myoblasts depleted of KEAP1, CCNF, or both to RNA-sequencing. As expected, lowering KEAP1 increased the mRNA levels of several NRF2 targets (**Figure 1C**). By contrast, depleting CCNF altered the expression of cell cycle regulators, likely due to the role of SCF^CCNF^ in degrading E2F transcription factors ^48^. Cells depleted of both KEAP1 and CCNF showed the expression profile caused by CCNF reduction and did not induce NRF2 targets (**Figure 1C**). qPCR analyses confirmed that loss of CCNF blunted the expression of NRF2 targets when KEAP1 levels were low (**Figure 1D**), an observation that we also made when KEAP1 was inhibited by chemical inducers of oxidative stress (**Figure 1E; Figure S1C**). These results were mirrored by changes in ROS: while depletion of KEAP1 reduced ROS levels dependent on NRF2 (**Figure S1D**), lowering CCNF had the opposite phenotype and re-equilibrated ROS levels in the absence of KEAP1 (**Figure S1E**). CCNF is therefore required for NRF2 activation and ROS scavenging even if KEAP1 levels are low and CUL3^KEAP1^ should not be fully active.

Having seen the effects of CCNF depletion onto NRF2 activity, we asked whether CCNF is important for myoblast survival during oxidative stress. Thus, we established a cell competition assay and mixed equal numbers of GFP-labeled control cells with mCherry-labeled cells depleted of KEAP1. After adding oxidative stressors, such as the glutathione synthesis inhibitor buthionine sulfoximine (BSO), we measured the ratio of GFP- to mCherry-labeled cells by flow cytometry. As expected, depleting KEAP1 improved myoblast survival dependent on NRF2 (**Figure S1F**). The protective effect of reducing KEAP1 was particularly apparent if cells experienced strong oxidative stress. Thus, some CUL3^KEAP1^ remains active even in the presence of high ROS, and this compromises the ability of myoblasts to survive such challenging conditions. Importantly, lowering CCNF eliminated the fitness benefit provided by KEAP1 inhibition during both moderate and strong oxidative stress (**Figure 1F**).

Together, these experiments showed that CCNF is required for oxidative stress signaling in myoblasts. CCNF is best known for targeting cell cycle regulators for proteasomal degradation ^48–51^. In addition, mutations in *CCNF* cause Amyotrophic Lateral Sclerosis (ALS) and Frontotemporal Dementia (FTD) ^52–55^, two diseases that affect non-dividing neurons and are characterized by ROS accumulation ^56^. It is possible that defective oxidative stress signaling, as observed here, contributes to the emergence of ALS or FTD in patients with *CCNF* mutations.

### p62/SQSTM supports oxidative stress signaling

As mutations in *CCNF* cause ALS, we asked if we could use myoblast differentiation to link other neurodegeneration risk factors to oxidative stress signaling. We therefore depleted proteins mutated in ALS together with KEAP1 and monitored myotube formation by microscopy. While CCNF-depletion showed the strongest phenotype, reducing p62 also allowed for substantial myogenesis despite low KEAP1 (**Figure 2A**). We confirmed with an independent siRNA that lowering p62 restored myotube formation in the absence of KEAP1 (**Figure 2B**). Loss of p62 limited the expression of some, but not all NRF2 targets, which is in line with its milder differentiation phenotype (**Figure 2C**), and depletion of p62 accordingly reversed the reduction in ROS levels that is observed upon NRF2 stabilization (**Figure 2D**). In line with these findings, p62 was required for the survival benefit of KEAP1 depletion during oxidative stress (**Figure 2E**). p62 therefore also supports robust oxidative stress signaling in myoblasts.

**Figure 2:**
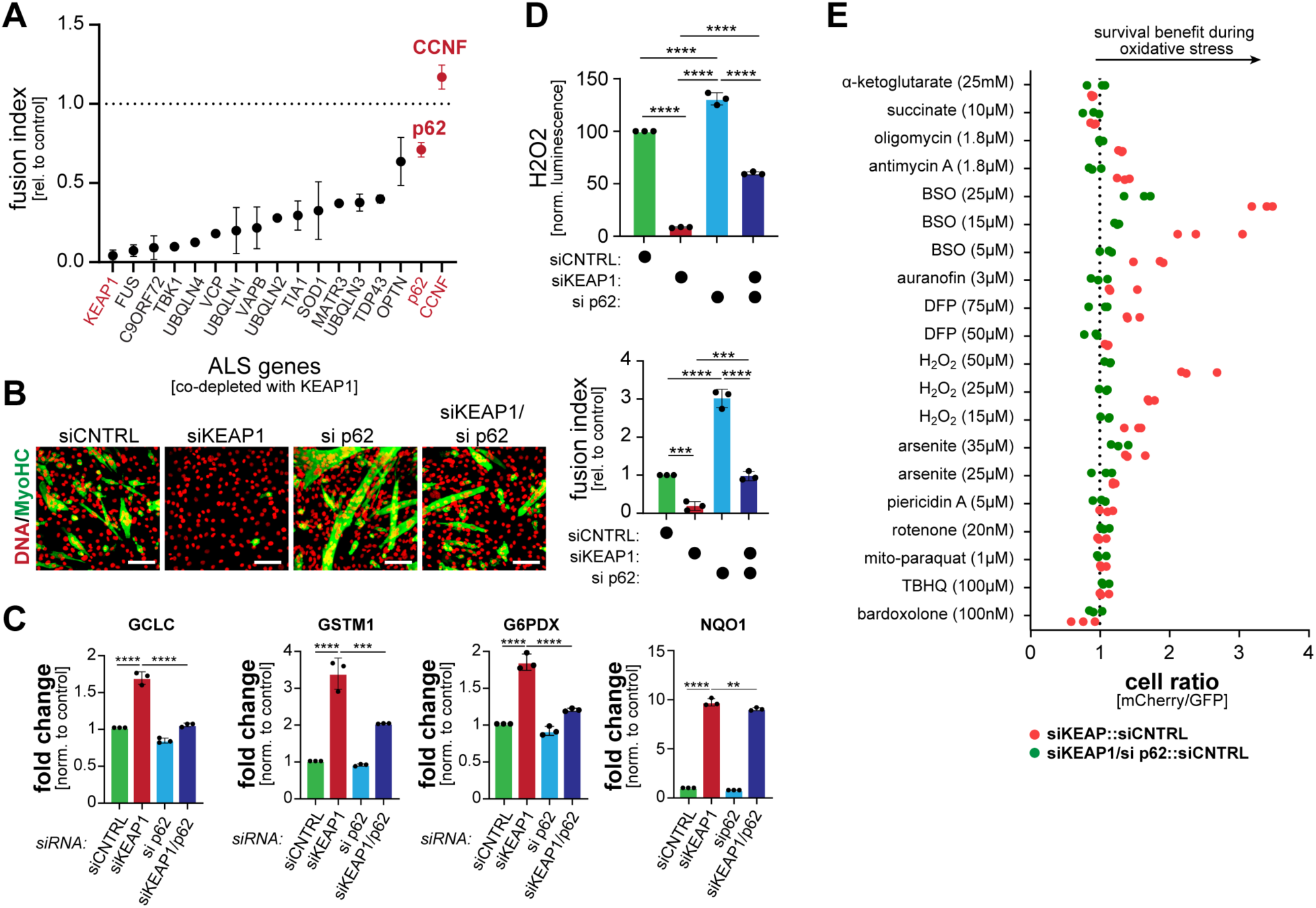
p62 supports oxidative stress signaling in myoblasts. **A.** C2C12 myoblasts were depleted of both KEAP1 as well as proteins encoding risk factors for Amyotrophic Lateral Sclerosis and Frontotemporal Dementia. After differentiation had been initiated, the success of myotube formation was monitored by immunofluorescence microscopy against MyoHC. Two independent screens were quantified. **B.** C2C12 myoblasts were depleted of KEAP1, p62, or both and induced to differentiate. Myotube formation was monitored by immunofluorescence microscopy against MyoHC. *Left:* microscopy images of differentiation; scale bar: 100μm. *Right:* quantification of three independent experiments. **C.** C2C12 myoblasts were depleted of KEAP1, p62, or both, and expression of NRF2 targets was determined by qPCR. n=3 replicates. **D.** C2C12 myoblasts were depleted of KEAP1, p62, or both, and intracellular ROS levels were determined by a ROS-Glo™ luciferase assay. **E.** GFP-labeled C2C12 myoblasts were mixed with mCherry-labeled cells that had been depleted of KEAP1 (red dots) or both p62 and KEAP1 (green dots). Cells were exposed to increasing concentrations of oxidative stressors. After two days, the ratio of GFP- to mCherry-labeled cells was determined by flow cytometry. Three independent experiments are shown (KEAP1-depleted cells are the same as in Figure 1F).

### NRF2 and CUL3^KEAP1^ bind the E3 ligase TRIP12

As a first step towards understanding how CCNF and p62 impact the oxidative stress response, we analyzed NRF2 levels in myoblasts depleted of KEAP1, CCNF, p62, or combinations thereof. While targeting KEAP1 led to the expected increase in NRF2, co-depletion of CCNF had a striking effect and prevented NRF2 accumulation (**Figure 3A**). Loss of CCNF blocked the increase in the NRF2 protein without impacting mRNA abundance (**Figure S2A-C**). p62 depletion also reduced NRF2 levels in cells with low KEAP1, but consistent with its milder differentiation phenotype, did so less drastically than loss of CCNF (**Figure 3B**). Treatment of cells co-depleted of CCNF and KEAP1 with the proteasome inhibitor carfilzomib, but not the lysosome inhibitor bafilomycin A, restored NRF2 (**Figure 3C**). These experiments showed that depletion of CCNF allowed myoblasts to target NRF2 for proteasomal degradation even if KEAP1 levels had been low.

**Figure 3:**
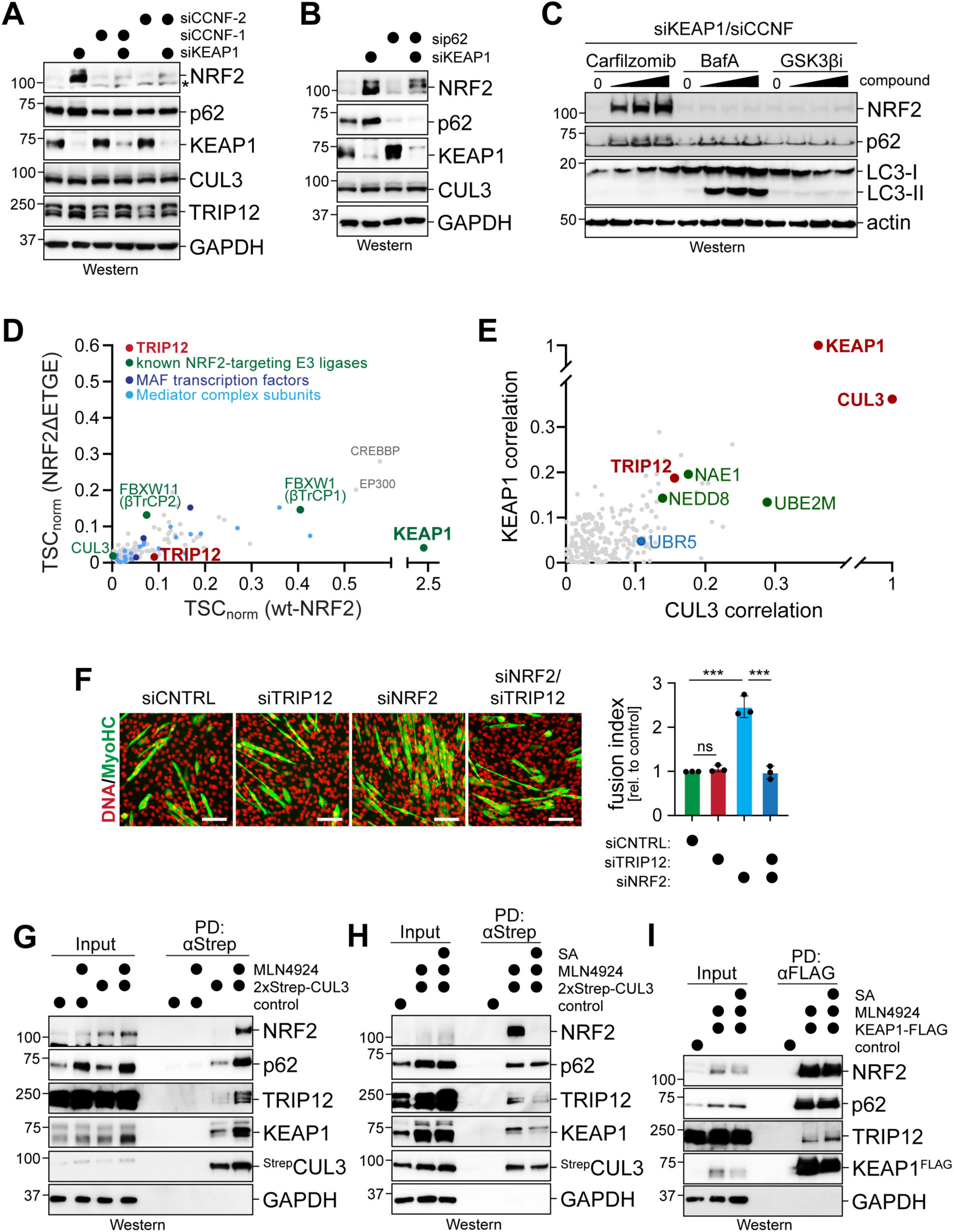
NRF2 and CUL3^KEAP1^ bind the E3 ligase TRIP12. **A.** C2C12 myoblasts were depleted of KEAP1, CCNF, or both, and NRF2 levels were determined by Western blotting. **B.** C2C12 myoblasts were depleted of KEAP1, p62, or both, and NRF2 levels were determined by Western blotting. **C.** C2C12 myoblasts were depleted of KEAP1 and CCNF at the same time and treated with increasing concentrations of the proteasome inhibitor carfilzomib; lysosome inhibitor bafilomycin A (BafA) or the GSK3β-inhibitor CHIR98014. NRF2 levels were determined by Western blotting. **D.** ^FLAG^NRF2 or ^FLAG^NRF2^ΔETGE^ were affinity-purified from lysates of stably transduced 293T cells that had been treated with MLN4924. Binding partners of NRF2 were determined by CompPASS mass spectrometry ^95^. **E.** DepMap analyses reveal TRIP12 as one of the most strongly correlated E3 ligases with CUL3 and KEAP1. Shown are all positive correlations of E3 ligases with either KEAP1 or CUL3, as derived from DepMap. **F.** C2C12 myoblasts were depleted of NRF2, TRIP12, or both. Following differentiation, the success of myotube formation was monitored by immunofluorescence against MyoHC. Quantification of three independent experiments is shown on the right. **G.** ^Strep^CUL3 was stably expressed in C2C12 myoblasts, affinity-purified, and analyzed for bound proteins by Western blotting using specific antibodies. As indicated, MLN4924 was added to delay substrate release from CUL E3 ligases. **H.** Stably expressed ^Strep^CUL3 was affinity-purified from C2C12 myoblasts, as described above. As indicated, sodium arsenite (25μM; 12h) was added to induce oxidative stress. MLN4924 was used to prevent substrate ubiquitylation and degradation. Bound proteins were detected by Western blotting using specific antibodies. **I.** ^FLAG^KEAP1 was transiently expressed in C2C12 myoblasts, affinity-purified and analyzed for binding partners by Western blotting using specific antibodies. As indicated, sodium arsenite (25μM; 12h) was added to induce oxidative stress.

How CCNF affects NRF2 stability is currently unknown and may involve indirect effects due its role in cell cycle regulation ^57^. In addition to revealing an unexpected function for CCNF, our observations also suggested that myoblasts possess mechanisms to sustain NRF2 degradation when CUL3^KEAP1^ had been compromised. This could include E3 ligases that boost CUL3^KEAP1^ activity and hence ensure dynamic regulation of NRF2 degradation. Previous work had implicated the E3 ligase SCF^βTrCP^ in targeting NRF2 that had been phosphorylated by GSK3 ^58^, but GSK3 inhibition by CHIR98014 did not stabilize NRF2 in the absence of CCNF and KEAP1 (**Figure 3C**). We therefore hypothesized that distinct ubiquitylation enzymes can support NRF2 turnover, if CUL3^KEAP1^ is not fully active.

To find such E3 ligases, we searched for binding partners of NRF2 by affinity-purification and mass spectrometry, using cells in which we stabilized NRF2 through the NEDD8-E1 inhibitor MLN4924 (**Figure 3D; Table S1**). We assessed wildtype NRF2 as well as a mutant, NRF2^ΔETGE^, that is recognized less efficiently by KEAP1. The two major NRF2 E3 ligases, KEAP1 and βTrCP, emerged as abundant interactors of NRF2, and binding of KEAP1 was strongly reduced by mutation of the ETGE motif in NRF2. E3 ligases that were proposed to target NRF2 in different cell types, such as HRD1 or WDR23 ^59,60^, did not associate with NRF2 in our experiments. However, we noted that NRF2, but less so NRF2^ΔETGE^, associated with the E3 ligase TRIP12 (**Figure 3D; Table S1**). Indicating that this interaction is of functional relevance, TRIP12 is correlated with CUL3^KEAP1^ in DepMap to the same extent as the NEDD8-E1 that is required for Cullin-RING ligase activity (**Figure 3E**) ^61^. Moreover, reducing TRIP12 alleviated the effects of NRF2 depletion onto myogenesis (**Figure 3F**), as expected if TRIP12 modulated the stability of this transcription factor.

As TRIP12 interacted less efficiently with NRF2^ΔETGE^, it may engage NRF2, at least in part, via binding to CUL3^KEAP1^. To test this notion, we purified CUL3 from myoblasts that were treated with MLN4924 to inhibit CUL3-dependent ubiquitylation and delay substrate release. We found that TRIP12 bound CUL3, which was enhanced by MLN4924 (**Figure 3G**). If cells experienced oxidative stress, CUL3 lost its association with NRF2 and bound slightly less TRIP12 (**Figure 3H**). KEAP1 also co-precipitated TRIP12 (**Figure 3I**), which showed that TRIP12 engages both NRF2 and its established E3 ligase CUL3^KEAP1^.

TRIP12 is an essential HECT-family E3 ligase that is highly expressed in brain and muscle ^62–64^. Mutations in *TRIP12* cause Clark-Baraitser Syndrome, which is characterized by autism-like symptoms that have also been linked to *CUL3* mutations ^44–46,65,66^. TRIP12 is overexpressed in Parkinson’s Disease ^47^, a neurodegenerative pathology characterized by ROS accumulation ^56^. Somatic mutations in *TRIP12* were observed in lung adenocarcinoma ^67^, a tumor caused by *KEAP1* or *NFE2L2* mutations that lead to NRF2 stabilization ^22,68^. Together with its interaction with CUL3^KEAP1^, these observations suggested that TRIP12 could support NRF2 degradation to establish the dynamic regulation of the oxidative stress response.

### TRIP12 is a ubiquitin chain elongation factor for CUL3^KEAP1^

To test whether TRIP12 targets NRF2, we reconstituted the ubiquitylation of NRF2 using purified components. We first incubated NRF2 with its established E3 ligase, CUL3^KEAP1^, which led to the decoration of NRF2 with the expected ubiquitin polymers (**Figure 4A**). Surprisingly, when we performed this reaction with ubiquitin variants containing a single lysine, we found that CUL3^KEAP1^ possessed little linkage specificity, except that it failed to produce K29-linked conjugates (**Figure 4B**). Also with p62 as substrate, CUL3^KEAP1^ did not assemble ubiquitin polymers if K29 was the only available lysine (**Figure 4C**), and a linkage-specific ubiquitin antibody showed that CUL3^KEAP1^ did not synthesize K29-linkages on NRF2 using wildtype ubiquitin (**Figure S3A**). Conversely, CUL3^KEAP1^ was more efficient in polyubiquitylating NRF2, if K29 had been mutated (**Figure S3B**). CUL3^KEAP1^ therefore displays ‘linkage exclusivity’: it can use any ubiquitin lysine, except K29, for polymer formation. A consequence of this activity, every ubiquitin attached to NRF2 by CUL3^KEAP1^ has K29 available for further modification.

**Figure 4:**
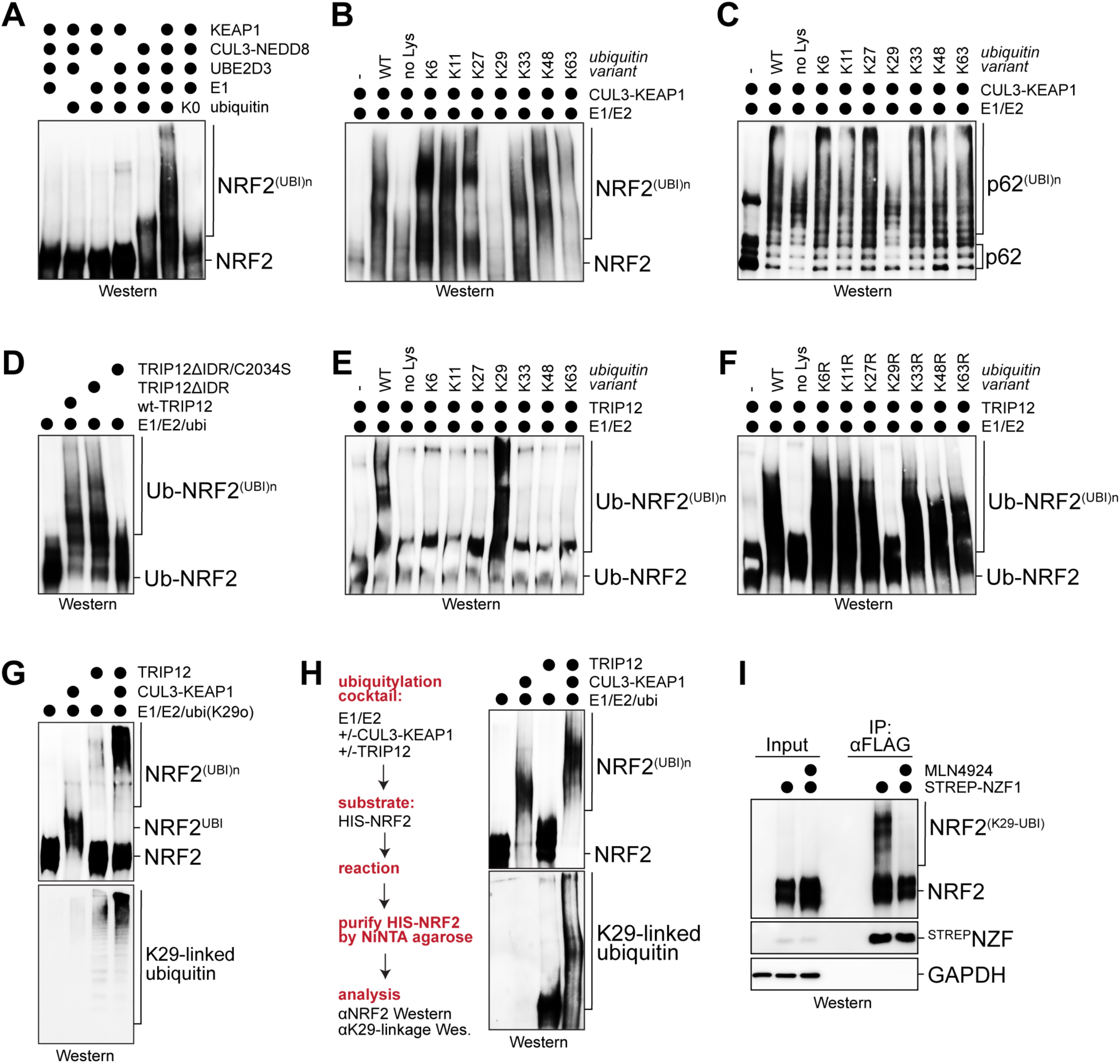
TRIP12 is a ubiquitin chain elongating factor for NRF2. **A.** Ubiquitylation of purified NRF2 by NEDD8-modified CUL3^KEAP1^ in the presence of wt-ubiquitin or a ubiquitin variant containing no Lys residues (ubi-K0). Reaction products were analyzed by Western blotting using αNRF2-antibodies. **B.** Ubiquitylation of NRF2 by CUL3^KEAP1^ in the presence of ubiquitin mutants that contained only a single Lys residue (K6: all Lys residues, except for K6, mutated to Arg). Reaction products were analyzed by Western blotting using αNRF2-antibodies. **C.** Ubiquitylation of purified p62 by CUL3^KEAP1^ in the presence of ubiquitin mutants that contained only a single Lys residue. Reaction products were analyzed by Western blotting using αp62-antibodies. **D.** Ubiquitylation of purified Ub∼NRF2 by TRIP12, a TRIP12 variant lacking its N-terminal intrinsically disordered region (TRIP12ΔIDR), or a TRIP12ΔIDR variant that also had its catalytic Cys residue mutated. Reaction products were analyzed by Western blotting using αNRF2-antibodies. **E.** Ubiquitylation of Ub∼NRF2 by TRIP12 in the presence of ubiquitin mutants containing a single Lys residue. Reaction products were analyzed by Western blotting using αNRF2-antibodies. **F.** Ubiquitylation of Ub∼NRF2 by TRIP12 in the presence of ubiquitin mutants that lacked a single Lys residue (K6R: only Lys6 of ubiquitin mutated to Arg). Reaction products were analyzed by Western blotting using αNRF2-antibodies. **G.** Ubiquitylation of NRF2 by CUL3^KEAP1^, TRIP12, or both, using a ubiquitin variant containing K29 as its only Lys residue. Reaction products were analyzed by Western blotting using antibodies against NRF2 and K29-linked ubiquitin chains. **H.** Ubiquitylation of purified ^HIS^NRF2 by CUL3^KEAP1^, TRIP12, or both in the presence of wt-ubiquitin. ^HIS^NRF2 was purified from reaction mixtures and analyzed for ubiquitylation by Western blotting using antibodies against NRF2 or K29-linked ubiquitin chains. **I.** NRF2 is modified in cells with ubiquitin conjugates containing abundant K29-linkages. C2C12 myoblasts expressed NRF2 as well as a Strep-tagged NZF1 domain of the K29/K33-specific DUB TRABID. ^STREP^NZF1 was affinity-purified from lysates under stringent conditions, and co-precipitating NRF2 was detected using αNRF2 antibodies. Where indicated, MLN4924 was added to prevent the chain initiation event required for TRIP12-dependent chain elongation.

TRIP12 did not modify NRF2 with ubiquitin chains of high molecular weight (MW) (**Figure S3C**). As TRIP12 had been linked to the ubiquitin-fusion pathway ^69,70^, we asked whether it instead extends ubiquitin conjugates that had already been attached to NRF2 by distinct enzymes, such as CUL3^KEAP1^. We therefore incubated TRIP12 with NRF2 that was fused to ubiquitin to bypass the initiation step (Ub∼NRF2) and found that TRIP12 efficiently modified this protein (**Figure 4D; Figure S3C**). The ability of TRIP12 to target Ub∼NRF2 required the catalytic Cys of the HECT domain, but not an intrinsically disordered region at its N-terminus (**Figure 4D**). Performing this experiment with ubiquitin variants revealed that K29 of ubiquitin was required and sufficient for modification of Ub∼NRF2 by TRIP12 (**Figure 4E, F**), and mutating K29 in the fused ubiquitin abolished TRIP12 activity towards Ub∼NRF2 (**Figure S3C**). As seen for other substrates ^71^, TRIP12 therefore extends K29-linked conjugates on Ub∼NRF2, a specificity that is complementary to the linkage exclusivity of CUL3^KEAP1^.

Based on these findings, we asked if CUL3 and TRIP12 cooperate to efficiently ubiquitylate NRF2. We incubated NRF2 with ubi-K29 and CUL3^KEAP1^ to allow for chain initiation, but not elongation, and then added TRIP12 to assess further modification. Reflecting their complementary linkage specificities, only the combination of CUL3^KEAP1^ and TRIP12 stimulated formation of high-MW conjugates, and K29-linkages were only detected when TRIP12 had been included in the reaction (**Figure 4G**). We next performed this experiment using wildtype ubiquitin and purified the modified NRF2 for analysis with linkage-specific antibodies ^72^. This strategy confirmed that CUL3^KEAP1^ did not produce any detectable K29-linked conjugates on NRF2, underscoring its intriguing linkage exclusivity (**Figure 4H**). While we noted that TRIP12 by itself can decorate NRF2 with K29-linked conjugates that did not change the mobility of NRF2 in gel electrophoresis, the combination of CUL3^KEAP1^ and TRIP12 resulted in polymers of higher MW, and K29-linkages were only detected if TRIP12 was included in this reaction. In line with these results, we found that NRF2 was modified in cells with conjugates that were enriched by a K29/K33-linkage specific ubiquitin-binding domain (**Figure 4I**). The K29-specific ubiquitylation of NRF2 was prevented by MLN4924, as expected due to its effect on inhibiting the first step of CUL3^KEAP1^/TRIP12-dependent modification. We conclude that TRIP12 is a ubiquitin chain elongation factor that cooperates with CUL3^KEAP1^ to ubiquitylate NRF2. TRIP12 adds K29-linked ubiquitin chains to NRF2, a chain topology that has been reported to drive efficient proteasomal degradation during stress ^71,73^.

### TRIP12 restrains NRF2 accumulation

TRIP12 is a predominantly nuclear protein ^74,75^. We therefore asked whether TRIP12 limits the accumulation of NRF2 in the nucleus, from which NRF2 is excluded by KEAP1-dependent sequestration and CUL3^KEAP1^-dependent degradation. Despite having weaker effects than lowering KEAP1, depletion of TRIP12 increased the abundance of nuclear NRF2 (**Figure 5A**). Targeting KEAP1 and TRIP12 at the same time led to further accumulation of nuclear NRF2 (**Figure 5A**). While this effect is likely due to incomplete depletion by siRNAs, it highlights that TRIP12 supports NRF2 degradation even if CUL3^KEAP1^ is not fully active.

**Figure 5:**
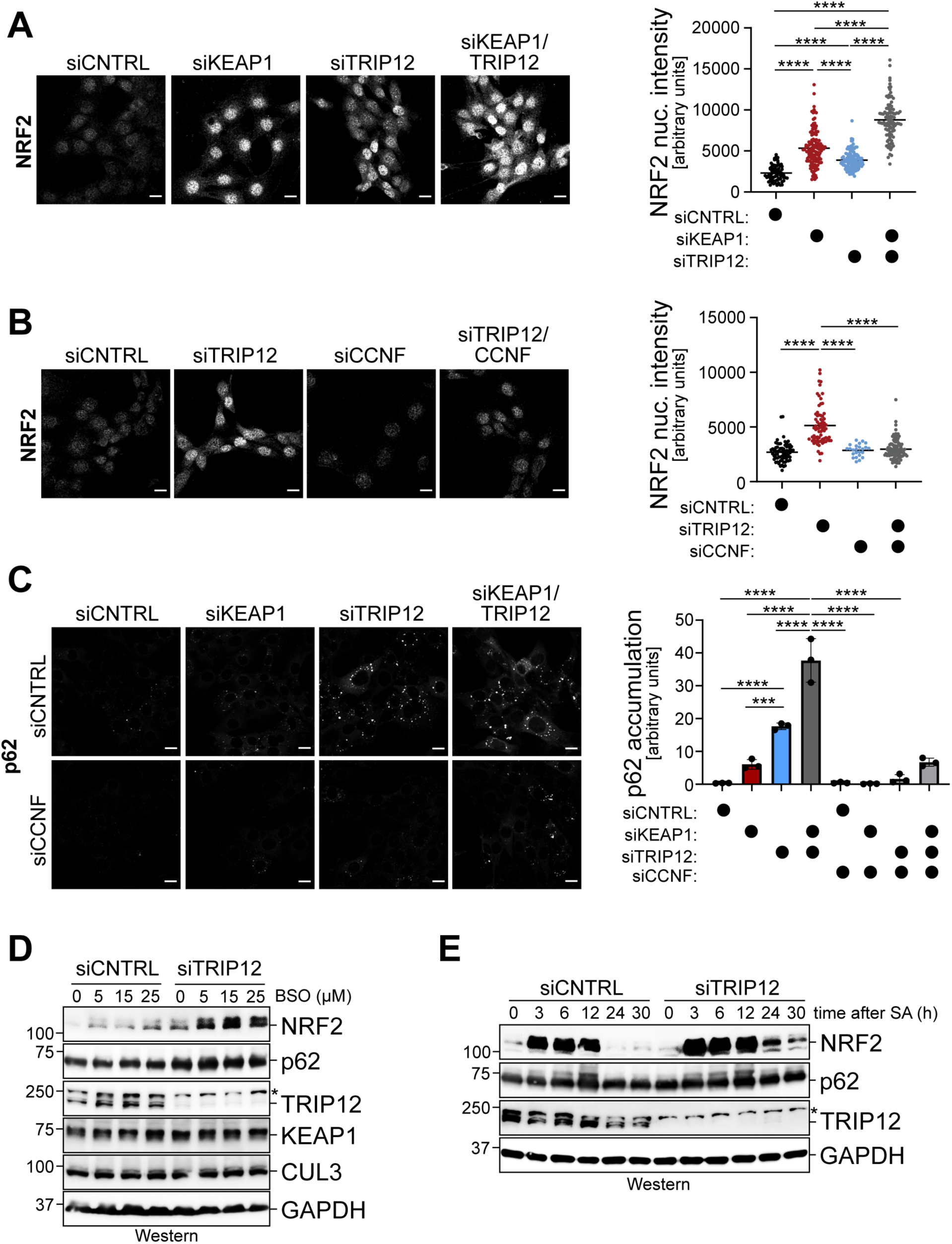
TRIP12 restricts accumulation of CUL3^KEAP1^ substrates. **A.** Depletion of KEAP1, TRIP12, or both in C2C12 myoblasts results in accumulation of nuclear NRF2, as determined by immunofluorescence against endogenous NRF2. The increase of nuclear NRF2 in cells treated with siRNAs against KEAP1 and TRIP12 at the same time is likely due to incomplete depletion of each target. Quantification is shown on the right. Scale bar: 10 μm. **B.** Accumulation of nuclear NRF2 in C2C12 myoblasts upon TRIP12 depletion is lost upon simultaneous inactivation of CCNF, as determined by microscopy against endogenous NRF2. Quantification is shown on the right. Scale bar: 10 μm. **C.** Depletion of KEAP1 or TRIP12 in C2C12 myoblasts results in accumulation of p62 punctae, as shown by microscopy against endogenous p62. Co-depletion of CCNF reverts effects of KEAP1- or TRIP12-loss. Quantification is shown on the right. Scale bar: 10 μm. **D.** Depletion of TRIP12 in C2C12 myoblasts further stabilizes NRF2 in cells exposed to increasing concentrations of the oxidative stressor BSO, as shown by Western blotting against endogenous proteins. **E.** Depletion of TRIP12 in C2C12 myoblasts delays degradation of NRF2 after recovery from sodium arsenite-induced oxidative stress (25μM for indicated times), as shown by Western blotting against endogenous proteins.

The consequences of TRIP12 depletion onto NRF2 accumulation are specific: neither loss of UBR5, which is correlated with TRIP12 on DepMap and known to target many transcription factors ^61,76^, nor depletion of other E3 ligases detected in some NRF2-immunoprecipitations affected the levels of NRF2 in the nucleus (**Figure S4A, B**). By contrast, lowering CCNF blunted the increase in nuclear NRF2 in TRIP12- or KEAP1-depleted cells (**Figure 5B**; **Figure S4C**), which suggests that TRIP12 and KEAP1 act in the same pathway to regulate NRF2. In line with these findings, depletion of TRIP12 reduced ROS levels in myoblasts, which was counteracted by concomitant loss of NRF2 or CCNF (**Figure S4D, E**).

In addition to stabilizing NRF2, CUL3^KEAP1^ inhibition caused the formation of cytoplasmic p62 foci (**Figure 5C**). Lowering TRIP12 also increased the abundance of p62 foci and, in fact, had a more pronounced effect than loss of KEAP1 (**Figure 5C**). As seen for NRF2, co-depleting KEAP1 and TRIP12 led to greater accumulation of p62 foci than either siRNA alone. Thus, TRIP12 also sustains regulation of p62, a protein that can shuttle between cytoplasm and nucleus ^77^. Loss of CCNF counteracted the effects of KEAP1- or TRIP12-depletion on the formation of p62 foci, which again underscores that KEAP1 and TRIP12 function in the same pathway. We conclude that TRIP12 regulates both NRF2 and p62, and this was particularly noticeable if KEAP1 levels were already low.

Rather than being depleted by siRNAs, CUL3^KEAP1^ is usually inhibited by ROS that accumulate during oxidative stress. To test if TRIP12 maintains NRF2 degradation during stress, we exposed myoblasts to increasing concentrations of BSO, hydrogen peroxide, or sodium arsenite. We then monitored NRF2 accumulation by Western blotting, which is less sensitive than microscopy but allowed us to compare more conditions at the same time. NRF2 levels increased in cells treated with BSO, peroxide, or sodium arsenite, and importantly, this effect was more pronounced when TRIP12 was depleted (**Figure 5D; Figure S4F, G**). TRIP12 therefore also sustains NRF2 turnover when CUL3^KEAP1^ activity has been downregulated during stress.

The continuous degradation of NRF2 during stress might allow cells to restore NRF2 turn-over more rapidly as ROS are being cleared. As arsenite elicits transient stress ^78^, we used this compound to test this hypothesis. In control cells, NRF2 accumulated upon exposure to arsenite and was degraded again as cells were recovering from stress (**Figure 5E**). By contrast, NRF2 persisted much longer in TRIP12-depleted cells (**Figure 5E**). Thus, TRIP12 supports NRF2 degradation as cells recover from oxidative stress and CUL3^KEAP1^ is being reactivated. We conclude that TRIP12 plays an important role in establishing the dynamic regulation of NRF2 degradation.

### TRIP12 restricts oxidative stress signaling

Having seen that TRIP12 restricts NRF2 accumulation, we wished to determine if it also impacts antioxidant gene expression. Partial inactivation of TRIP12, as accomplished by siRNAs, was not sufficient to elicit a pronounced spike in the expression of NRF2 targets (**Figure 6A-C**). In a similar manner, moderate concentrations of BSO, sodium arsenite, or oligomycin did not trigger a persistent oxidative stress response. However, if TRIP12-depleted cells were exposed to the same oxidative insults, many NRF2 targets were strongly induced (**Figure 6A-C**). Depletion of TRIP12 increased the expression of NRF2 targets that were relevant to the underlying biology: for example, BSO-dependent inhibition of glutathione synthesis in TRIP12-depleted cells resulted in strong upregulation of the GCLM subunit of glutamate-cysteine ligase, while arsenite-dependent induction of oxidative and proteotoxic stress led to pronounced expression of p62 as a regulator of autophagy (**Figure 6A-C**). These experiments showed that TRIP12 restricts NRF2 activation, which was noticeable if CUL3^KEAP1^ had been compromised by moderate oxidative stress.

**Figure 6:**
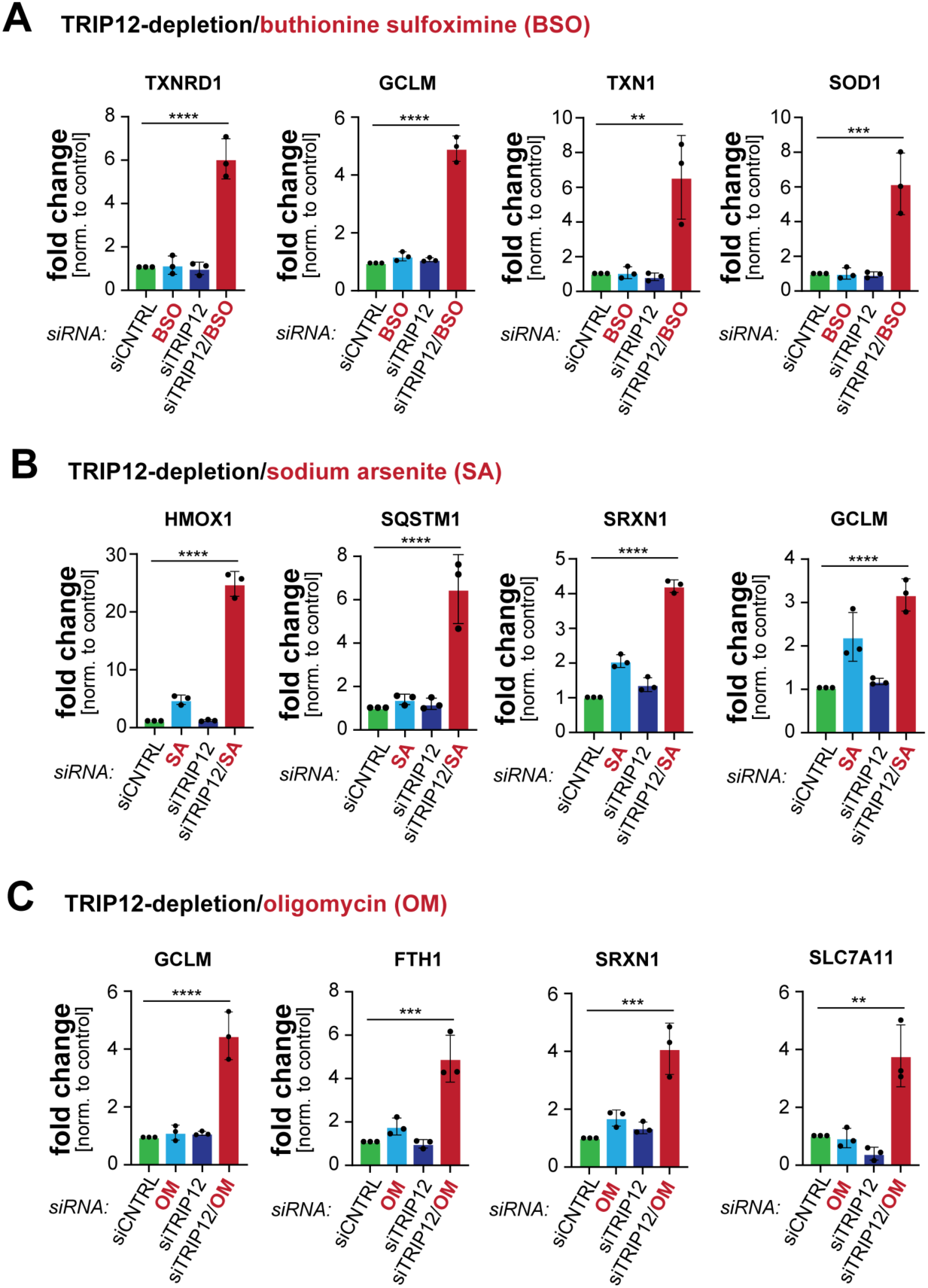
TRIP12 restricts antioxidant gene expression. **A.** C2C12 myoblasts were treated with the inhibitor of glutathione synthesis, BSO (16h; 5μM); depleted of TRIP12; or both. Expression of NRF2 target genes was determined by qPCR. n=3 replicates. **B.** C2C12 myoblasts were treated with sodium arsenite (16h; 5μM); depleted of TRIP12; or both. Expression of NRF2 target genes was determined by qPCR. n=3 replicates. **C.** C2C12 myoblasts were treated with the mitochondrial ATP synthase inhibitor oligomycin (16h; 0.9μM); depleted of TRIP12; or both. Expression of NRF2 target genes was determined by qPCR. n=3 replicates.

To investigate if TRIP12 helps to shut down antioxidant gene expression during recovery from stress, we treated cells with a pulse of sodium arsenite and monitored mRNA levels at times when NRF2 is usually being degraded again (see **Figure 5E**). While control cells accordingly shut down NRF2 targets at this time, cells depleted of TRIP12 continued to express several antioxidant proteins and hence showed a delay in silencing the oxidative stress response (**Figure 7A**). Thus, while TRIP12 limits NRF2 activation during stress, it accelerates the return of cells to homeostatic conditions by shutting off antioxidant gene expression.

**Figure 7:**
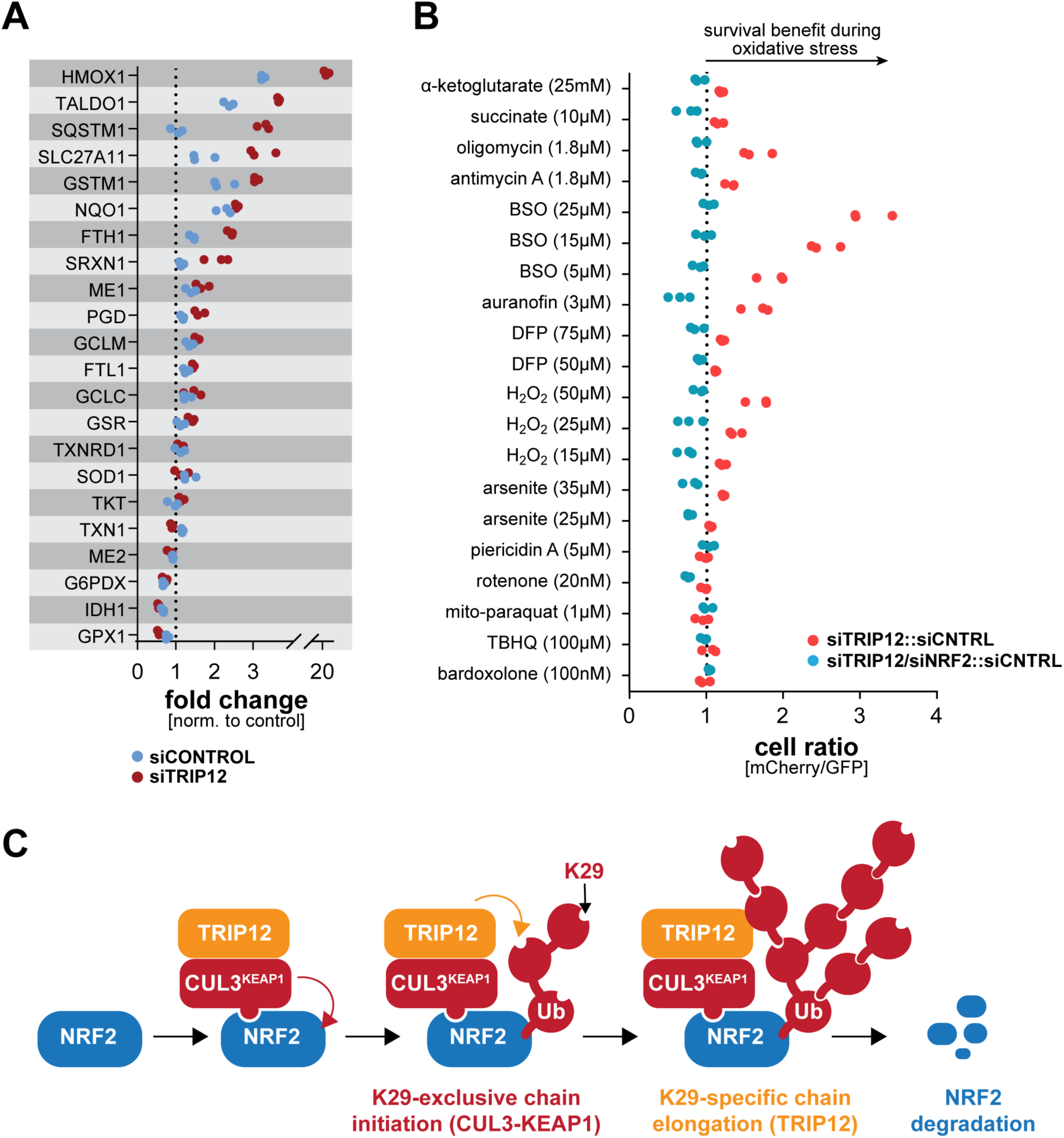
TRIP12 restricts oxidative stress signaling and cell survival during stress. **A.**C2C12 myoblasts were transfected with control siRNAs or siRNAs targeting TRIP12. Oxidative stress was induced by exposing cells to sodium arsenite (24h; 25μM). After 24h, when control cells had reactivated NRF2 degradation, expression of NRF2 target genes was determined by qPCR. **B.** GFP-labeled control cells were mixed at a 1:1 ratio with mCherry-labeled cells depleted of TRIP12 (red dots). As indicated, NRF2 was also depleted (blue dots). Cells were exposed to increasing concentrations of oxidative stressors. After two days, the ratio of GFP- to mCherrylabeled cells was determined by flow cytometry. n=3 independent experiments. **C.** Model of NRF2 ubiquitylation carried out by CUL3^KEAP1^ and TRIP12.

TRIP12 therefore restricts both NRF2 accumulation and function, prompting us to ask whether its inhibition would boost cell survival during oxidative stress. Strikingly, the depletion of TRIP12 had the same beneficial effect as reducing KEAP1 and improved myoblast survival upon glutathione synthesis inhibition by BSO; TXNRD1 inhibition by auranofin; ATPase inhibition by oligomycin; or exposure to H_2_O_2_ (**Figure 7B**). As seen for KEAP1, the protective effect of lowering TRIP12 was dependent on NRF2 (**Figure 7B**), and it required both CCNF and p62 (**Figure S5A, B**). We conclude that TRIP12 is a crucial component of the oxidative stress response that restricts NRF2 activation even during stress. This finding revealed that the need for dynamic control of the oxidative stress response comes at a cost: to ensure that ROS clearance is quickly followed by a return to homeostatic conditions, cells continue to degrade some NRF2 during stress despite this limiting their ability to survive challenging conditions.

## Discussion

Dynamic control of oxidative stress signaling is critical for cell and tissue homeostasis. When cells detect ROS, they rapidly stabilize NRF2 to induce antioxidant gene expression. However, as cells overcome oxidative insults, they must quickly eliminate NRF2 to prevent reductive stress and its deleterious consequence on cellular metabolism and function. Here, we show that dynamic regulation of NRF2 degradation and antioxidant signaling requires TRIP12, an E3 ligase mutated in the neurodevelopmental Clark-Baraitser Syndrome and overexpressed in Parkinson’s Disease. TRIP12 is a ubiquitin chain elongating factor that amplifies CUL3^KEAP1^-dependent conjugates to ensure robust NRF2 turnover (**Figure 7C**). In this manner, TRIP12 restricts NRF2 activation during stress, but facilitates NRF2 elimination during recovery. The ability of cells to ensure transient activation of the oxidative stress response is therefore so important that it comes at the cost of limited protective signaling in the face of ROS.

### Mechanism of NRF2 ubiquitylation

As illustrated by mutually exclusive cancer mutations in KEAP1 and NRF2 ^5^, CUL3^KEAP1^ has long been known to be a major E3 ligase for NRF2. However, several observations indicated that additional factors might boost NRF2 turnover, and we identify TRIP12 as one such protein. TRIP12 is a ubiquitin chain elongation factor, frequently referred to as E4 ^79^, which amplifies initial conjugates that were attached to NRF2 by CUL3^KEAP1^. While we discovered this role of TRIP12 in in myoblasts, DepMap analyses indicate that TRIP12 and CUL3^KEAP1^ work together in many other cell types. TRIP12 also functions in the ubiquitin-fusion pathway ^69,70^, where chain elongation factors were first described ^79^, and it collaborates with CUL1 and CUL4 E3 ligases to jumpstart targeted protein degradation ^71^. It therefore appears that TRIP12 amplifies ubiquitin signals of multiple E3 ligases. In line with being a central ubiquitylation enzyme, *TRIP12* is essential for mouse development ^63^, and its products, K29-linked ubiquitin conjugates, are abundant in human cells ^72^.

We identified TRIP12 by searching for binding partners of NRF2. TRIP12 interacted less efficiently with an NRF2 variant that lacked its major motif recognized by KEAP1, suggesting that TRIP12 is recruited to NRF2 via CUL3^KEAP1^. We indeed found that TRIP12 bound CUL3^KEAP1^, which was detected more prominently if ubiquitylation was inhibited and substrate release from CUL3^KEAP1^ was delayed. TRIP12 therefore appears to engage substrate-loaded CUL3^KEAP1^, but how this interaction occurs is unknown and will require structural analyses. Complex formation between TRIP12 and CUL3^KEAP1^ likely facilitates efficient ubiquitin handover from the initiation site in CUL3^KEAP1^ to the elongation module provided by TRIP12, thereby promoting robust NRF2 degradation (**Figure 7C**).

TRIP12 is well suited to cooperate with CUL3^KEAP1^, as biochemical experiments revealed that these E3 ligases possess surprising linkage complementarity. CUL3^KEAP1^ failed to modify K29 of ubiquitin, a catalytic preference that we refer to as K29-linkage exclusivity. Why CUL3^KEAP1^ does not target K29 of ubiquitin is unclear, but it might bind ubiquitin in a manner that orients this lysine away from a CUL3^KEAP1^-bound E2. The linkage exclusivity of CUL3^KEAP1^ results in conjugates in which every building block has K29 of ubiquitin available for chain elongation, and the responsible E3 ligase TRIP12 indeed has complementary K29-linkage specificity. We anticipate that TRIP12 acts on multiple ubiquitin subunits on each NRF2, thereby producing polymers that resemble branched ubiquitin chains for efficient proteasomal degradation ^80^. Branched chains recruit the segregase p97/VCP ^80–82^, which is required for NRF2 degradation ^83^. In DepMap, TRIP12 is correlated with the p97-interactors PLAA and VCPIP ^61^, and its products, K29-linked ubiquitin conjugates, co-localize with p97 in stressed cells ^72^. Together, these findings suggest that CUL3^KEAP1^ and TRIP12 together produce a ubiquitin signal that enables efficient NRF2 turnover even if cells experience stress or need to return to homeostatic conditions during recovery.

### Role of TRIP12-dependent ubiquitin chain extension

Our study identified TRIP12 as a negative regulator of NRF2 accumulation and function. Accordingly, TRIP12 depletion unleashed antioxidant gene expression if cells experienced moderate oxidative stress and CUL3^KEAP1^ has been partially inhibited. Inactivation of TRIP12 also delayed the silencing of NRF2-dependent transcription once ROS have been cleared, yet CUL3^KEAP1^ is not yet fully active. Thus, the TRIP12-dependent amplification of ubiquitin signals is particularly important when CUL3^KEAP1^ is less active and fewer ubiquitin chain initiation events are expected to occur. How could adding more ubiquitin molecules to NRF2 improve proteasomal degradation? We propose that without much linkage-specificity, the ubiquitin chains produced by CUL3^KEAP1^ are poor proteasomal targeting signals. TRIP12-dependent attachment of K29-linked conjugates adds defined ubiquitin blocks that improve the recognition of NRF2 by p97 and the proteasome to increase the efficiency of NRF2 degradation. A flipside of this regulation, if chain initiation and elongation are coordinated less efficiently, proteins bound to CUL3^KEAP1^ might escape proteasomal degradation.

Depletion of TRIP12 by siRNAs had weaker effects on NRF2 accumulation and antioxidant gene expression than lowering KEAP1. Thus, CUL3^KEAP1^ might control a rate-limiting step in NRF2 ubiquitylation and TRIP12 only becomes critical during times of stress or when cells need to return to homeostatic conditions. Alternatively, additional chain elongation factors might support some ubiquitin chain formation when TRIP12 had been depleted. However, it is important to note that we could not generate *ΔTRIP12* myoblasts and hence had to rely on siRNAs that partially deplete their target. While our failure to obtain *ΔTRIP12* myoblasts is consistent with TRIP12’s essential role in embryogenesis, it raises the possibility that TRIP12 is highly active and any remaining TRIP12 in siRNA-treated cells is sufficient to accomplish sufficient NRF2 ubiquitylation.

By depleting KEAP1 or TRIP12, we noticed that cells continue to degrade some NRF2 even if they experience strong oxidative stress. While the residual NRF2 turnover restricts protective signaling and limits cell survival, it eases the return of cells to homeostatic conditions as ROS are being cleared. Cells therefore establish transient stress response activation through a mechanism that reduces their ability to cope with stress. A failure to shut off NRF2 depletes ROS that are required for cell differentiation and tissue homeostasis ^14,15,17,84–87^. Recent work showed that such ROS are sentinel molecules that allow cells to finetune the activity of their electron transport chain, a pathway critical for producing the energy and metabolic building blocks that fuel cell fate decisions ^88^. Moreover, prolonged KEAP1 inhibition results in a form of cell death referred to as oxeiptosis ^20^, and inactivation of CUL3^KEAP1^ by mutations in *KEAP1* or *CUL3* drives multiple cancers ^7,22,89^. The importance of establishing transient stress response activation has been documented beyond CUL3^KEAP1^ and NRF2, such as for mitochondrial and proteotoxic stress responses ^78,90,91^. Together with the latter findings, our discovery of TRIP12 as a negative regulator of the oxidative stress response therefore underscores that dynamic regulation of stress signaling is critical for preserving organismal health.

The ability of cells to degrade NRF2 if CUL3^KEAP1^ has been inhibited became apparent upon depletion of the SCF/CUL1-adaptor CCNF. Lowering CCNF in myoblasts stimulated proteasomal turnover of NRF2 if KEAP1 was depleted, which prevented antioxidant gene expression and restored the ability of these cells to differentiate ^15^. How CCNF impacts NRF2 stability is unknown, but RNA-sequencing showed that CCNF depletion rewires gene expression consistent with previously reported effects on cell cycle progression ^48,50,51,57^. During the G1 phase of the cell cycle, the deubiquitylase USP11 is degraded to allow for CUL3^KEAP1^-dependent inhibition of homologous recombination ^35^. CCNF might similarly affect NRF2 turnover by changing the cell cycle stage and altering the abundance of counteracting deubiquitylases. Dissecting the mechanism through which CCNF impacts NRF2 stability is bound to uncover further regulation of oxidative stress signaling and could reveal why *CCNF* mutations cause neurodegenerative diseases that are characterized by aberrant ROS accumulation ^52,53,92^.

### Therapeutic potential of TRIP12

As ROS accumulation is a hallmark of many neurodegenerative diseases ^1,93^, transient NRF2 stabilization by small molecules might be of therapeutic benefit. While KEAP1 inhibition would be one approach to induce NRF2, it is dangerous: mutations in *KEAP1* or in the KEAP1-binding motif of NRF2 cause several cancers ^89^. *TRIP12* variants have also been observed in cancer, but are unlikely to be driver mutations. We suggest that TRIP12 inhibition provides a safer route to activating NRF2 without promoting tumorigenesis. At least in myoblasts, TRIP12 inhibition offers the same survival benefit during stress as loss of KEAP1 but preferentially increases NRF2 levels if cells already experience some ROS, as expected for neurons at the brink of degeneration. Our work therefore reveals a component of the oxidative stress response, TRIP12, that could be targeted to rewire this protective signaling system for therapeutic benefit.

### Limitations of the Study

While our study deepens the understanding of the oxidative stress response, a pathway linked to cancer and neurodegeneration, it raises several questions. How CCNF modulate NRF2 stability is unknown: does CCNF regulate the CUL3^KEAP1^-TRIP12 machinery, or does it act through an unknown E3 ligase? It is possible that CCNF’s function towards NRF2 is connected to its role as a cell cycle regulator, in line with observations that ROS impinge on the activity of cell cycle kinases ^94^. A technical limitation of our work, *TRIP12* is essential for embryogenesis and we failed to delete it from myoblasts. Thus, we had to rely on depletion of TRIP12 by siRNAs, rather than complete inactivation. It is therefore unclear whether TRIP12 is the only chain elongating factor for NRF2 or whether additional ubiquitylation enzymes help to establish dynamic regulation of the oxidative stress response. Finally, it will be important to determine whether other CUL3 E3 ligases, such as CUL3^SPOP^, rely on chain elongating enzymes that may include TRIP12. Given the prominent roles of CUL3^KEAP1^ or CUL3^SPOP^ in tumorigenesis, dissecting mechanisms of ubiquitin chain formation by these enzymes is an important area of study that holds promise for developing novel therapeutic approaches.

## Resource Availability

### Lead Contact

Further information and requests for reagents and resources should be directed to Michael Rapé.

### Materials Availability

All plasmids generated in this work can be requested from the lead contact’s lab and will be freely shared. All antibodies, chemicals, and cell lines used in this study are commercially available.

### Data Availability

Original proteomics data will be uploaded as a supplemental table. Original RNA sequencing data has been deposited into the NCBI gene expression omnibus (GSE281184O) and will be made publicly available upon acceptance. Additional information to reanalyze data reported in this work paper is available from the Lead Contact upon request.

## Acknowledgements

We are grateful to Diane Haakonsen, Rumi Sherriff, Zhi Yang and David Ye for thoughtful discussions. We thank all of our lab members for help and support. We are grateful to the CRL Flow Cytometry Facility, Vincent J. Coates Proteomics/Mass Spectrometry Laboratory (NIH S10RR025622) and HTS Facility (NIH S10OD021828) at UCB. AJI is a recipient of a National Science Foundation pre-doctoral fellowship. This work was supported by an NIGMS RO1 to MR (GM151335-01). MR is an HHMI investigator.

## Author contributions

AJI, DMM, and JYH all performed experiments. All authors analyzed the data. MR supervised the work. AJI and MR wrote the manuscript.

## Inclusion and Diversity

We support inclusive, diverse, and equitable conduct of research, and one or more of our authors identifies as an underrepresented minority in science.

## Supplementary Figures

**Figure S1, related to Figure 1:**
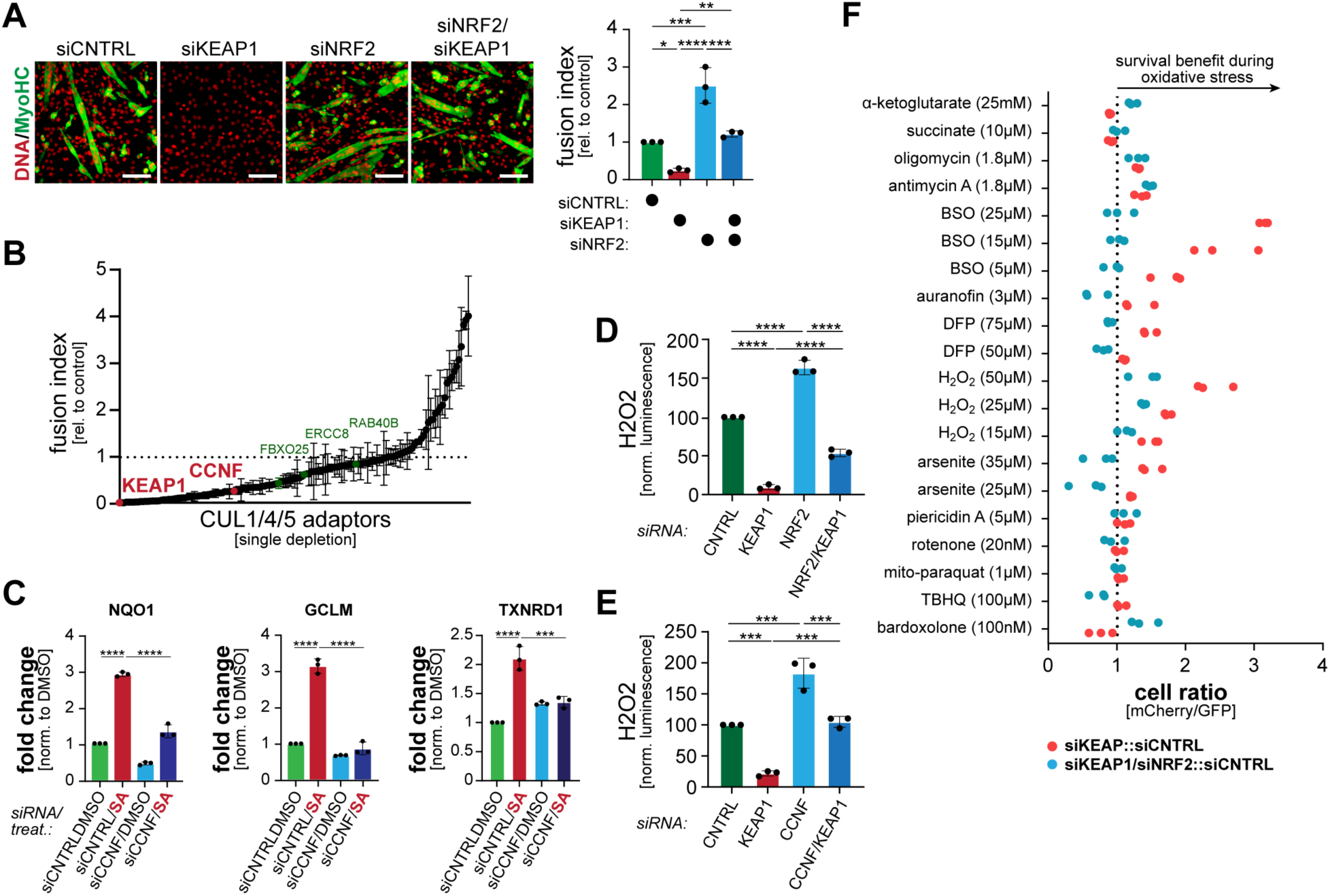
CCNF modulates oxidative stress signaling in myoblasts. **A.** C2C12 myoblasts were depleted of KEAP1, NRF2, or both. After differentiation, myotube formation was analyzed by immunofluorescence against MyoHC (green). Nuclei were stained with Hoechst (red). Quantification of three independent experiments is shown on the right. **B.** Substrate adaptors of CUL1, CUL4, and CUL5 E3 ligases were depleted from C2C12 myoblasts. After differentiation was initiated, the success of myotube formation was monitored by immunofluorescence microscopy against MyoHC. n=2 replicates. **C.** C2C12 myoblasts were transfected with control siRNAs or siRNAs against CCNF and exposed to sodium arsenite (16h; 25μM). Expression of select NRF2 targets was determined by qPCR. n=3 replicates. **D.** C2C12 myoblasts were depleted of KEAP1, NRF2, or both, and intracellular ROS were determined by a ROS-Glo™ H_2_O_2_ assay (Promega). **E.** C2C12 myoblasts were depleted of KEAP1, CCNF, or both, and intracellular ROS were determined by a ROS-Glo™ H_2_O_2_ assay (Promega). **F.** GFP-labeled control cells were mixed at a 1:1 ratio with mCherry-labeled cells depleted of KEAP1 (red dots). As indicated, NRF2 was also depleted (blue dots). Cells were exposed to increasing concentrations of oxidative stressors. After two days, the ratio of GFP- to mCherry-labeled cells was determined by flow cytometry. n=3 independent experiments. KEAP1-depleted cells are the same as shown in Figure 1F.

**Figure S2, related to Figure 3:**
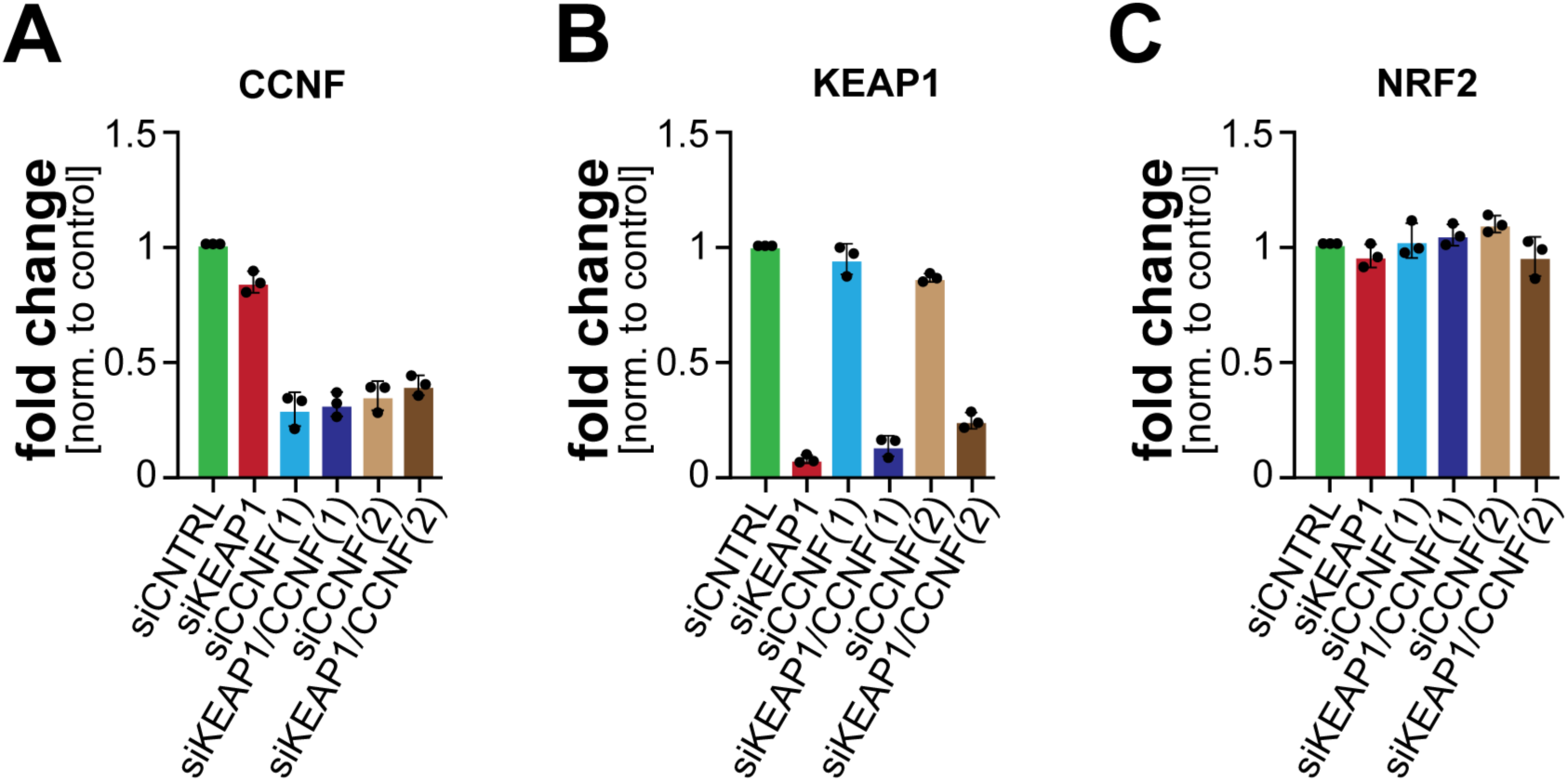
Depletion of CCNF does not affect NRF2 mRNA. **A.** C2C12 myoblasts were depleted of KEAP1, CCNF (2 independent siRNAs), or both, and the abundance of CCNF mRNA was determined by qPCR. **B.** C2C12 myoblasts were depleted of KEAP1, CCNF (2 independent siRNAs), or both, and the abundance of KEAP1 mRNA was determined by qPCR. **C.** C2C12 myoblasts were depleted of KEAP1, CCNF (2 independent siRNAs), or both, and the abundance of NRF2 mRNA was determined by qPCR.

**Figure S3, related to Figure 4:**
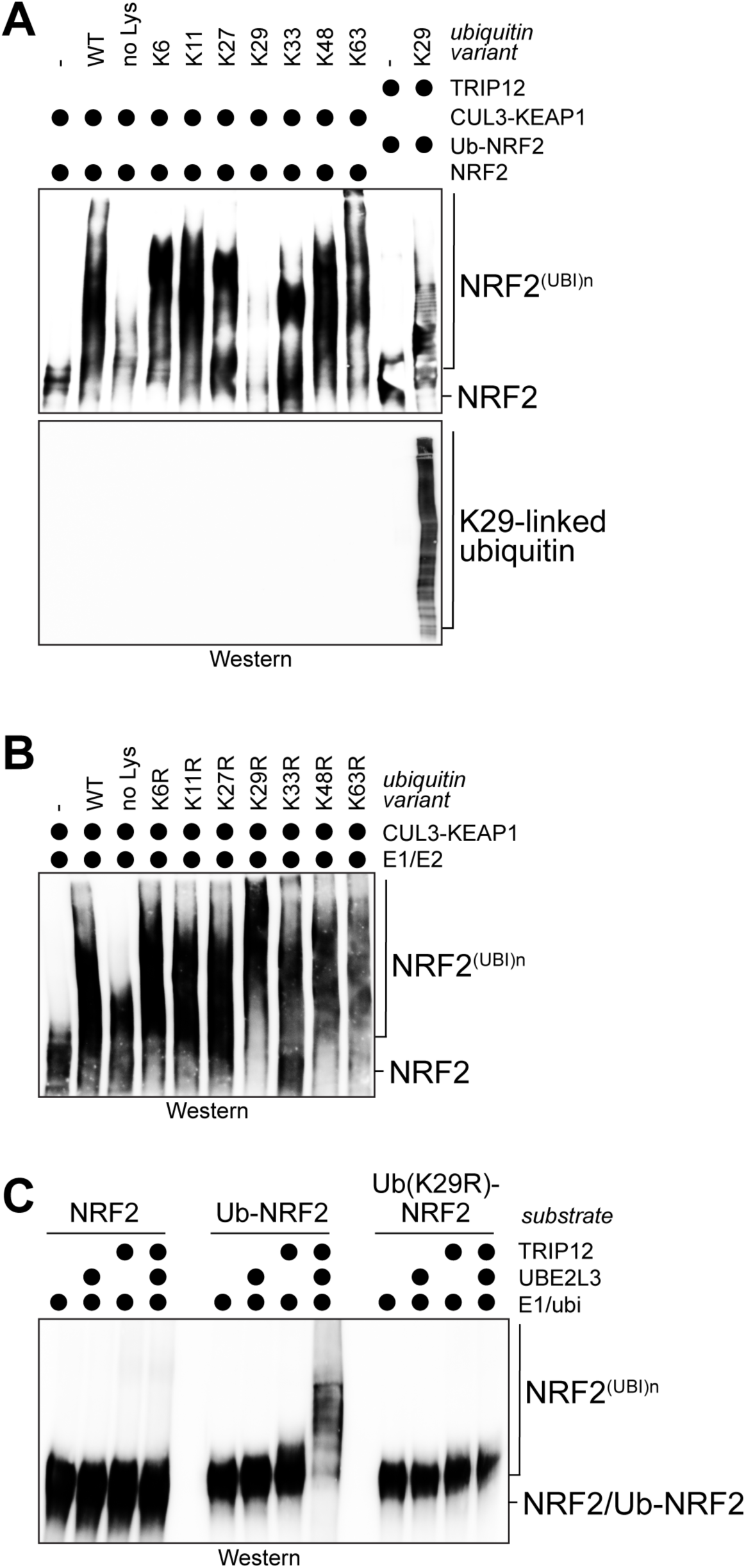
TRIP12 is a ubiquitin chain elongation factor for NRF2. **A.** CUL3^KEAP1^ does not assemble ubiquitin conjugates containing K29-linkages. Purified NRF2 was incubated with CUL3^KEAP1^, E1, UBE2D3, and either wildtype or single-Lys ubiquitin mutants. In the last two lanes, purified Ub∼NRF2 was incubated with TRIP12 and a ubiquitin variant containing Lys29 (positive control for K29-linkage formation). Reaction products were analyzed by Western blotting using antibodies against NRF2 or K29-linked ubiquitin chains. **B.** CUL3^KEAP1^-dependent ubiquitylation of NRF2 occurs more efficiently in the presence of a ubiquitin variant lacking K29 (ubi-K29R). Ubiquitylation of purified NRF2 was performed as described above. **C.** TRIP12^ΔIDR^ was purified from 293T cells and incubated with E1, UBE2L3 as a HECT-specific E2, ubiquitin, and either recombinant NRF2, Ub∼NRF2, or Ub^K29R^∼NRF2. Reaction products were analyzed by Western blotting against NRF2.

**Figure S4, related to Figure 5:**
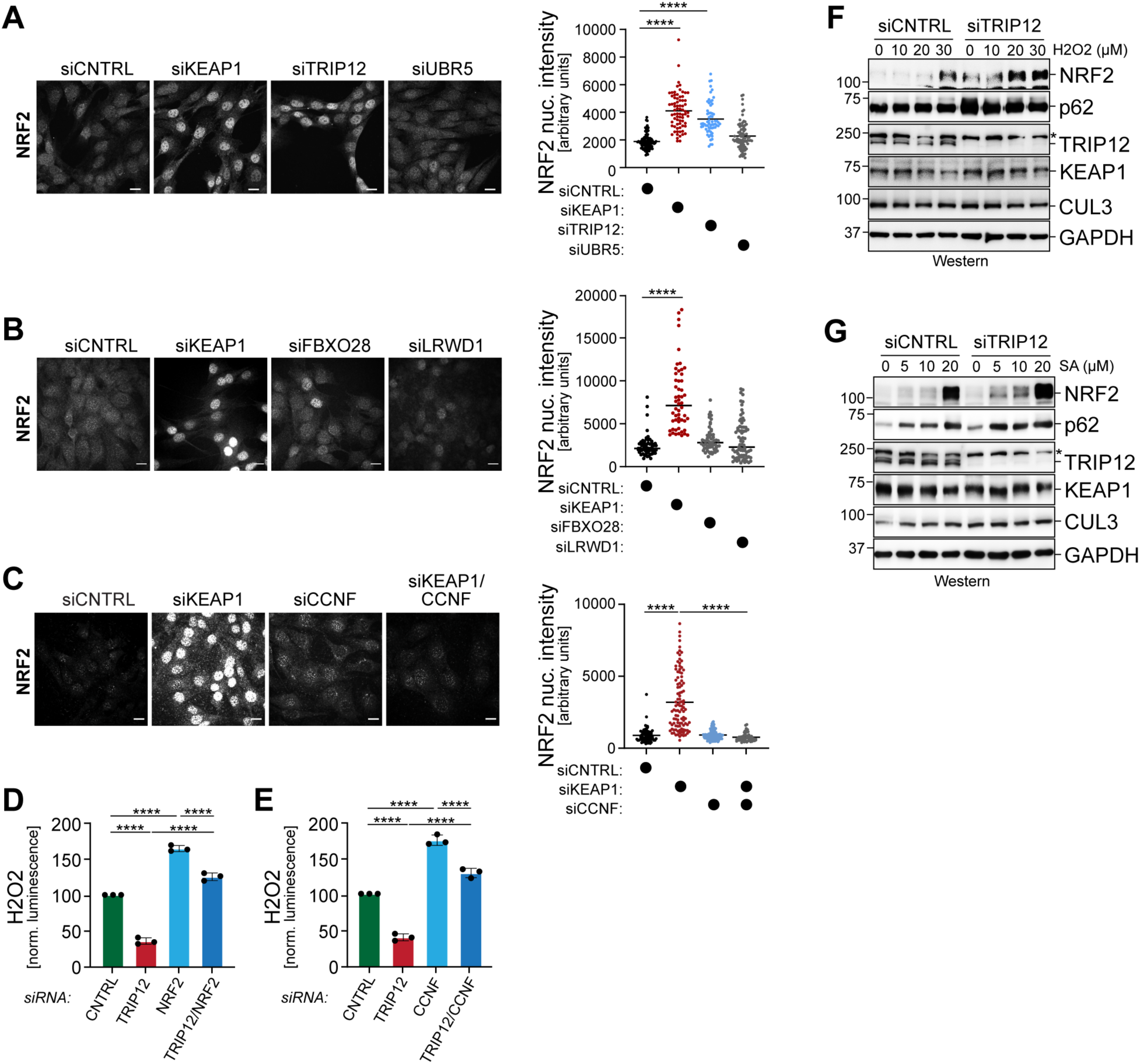
TRIP12 restricts NRF2 accumulation. **A.** C2C12 myoblasts were depleted of KEAP1, TRIP12, or UBR5, and levels of nuclear NRF2 were determined by immunofluorescence microscopy against endogenous NRF2. Quantification is shown on the right. UBR5 is the E3 ligase that is most closely correlated with TRIP12 across DepMap. **B.** C2C12 myoblasts were depleted of KEAP1, FBXO28, or LRWD1, and levels of nuclear NRF2 were determined by immunofluorescence microscopy against endogenous NRF2. Quantification is shown on the right. FBXO28 and LRWD1 were detected in NRF2 affinity-purification and mass spectrometry. **C.** C2C12 myoblasts were depleted of KEAP1, CCNF, or both, and levels of nuclear NRF2 were determined by immunofluorescence microscopy against endogenous NRF2. Quantification is shown on the right. **D.** C2C12 myoblasts were depleted of TRIP12, NRF2, or both, and intracellular ROS were determined by a ROS-Glo™ H_2_O_2_ assay (Promega). **E.** C2C12 myoblasts were depleted of TRIP12, CCNF, or both, and intracellular ROS were determined by a ROS-Glo™ H_2_O_2_ assay (Promega). **F.** C2C12 myoblasts were depleted of TRIP12 and exposed to increasing concentrations of hydrogen peroxide. Levels of NRF2 and additional proteins were determined by Western blotting using specific antibodies. **G.** C2C12 myoblasts were depleted of TRIP12 and exposed to increasing concentrations of sodium arsenite. Levels of NRF2 and additional proteins were determined by Western blotting using specific antibodies.

**Figure S5, related to Figure 7:**
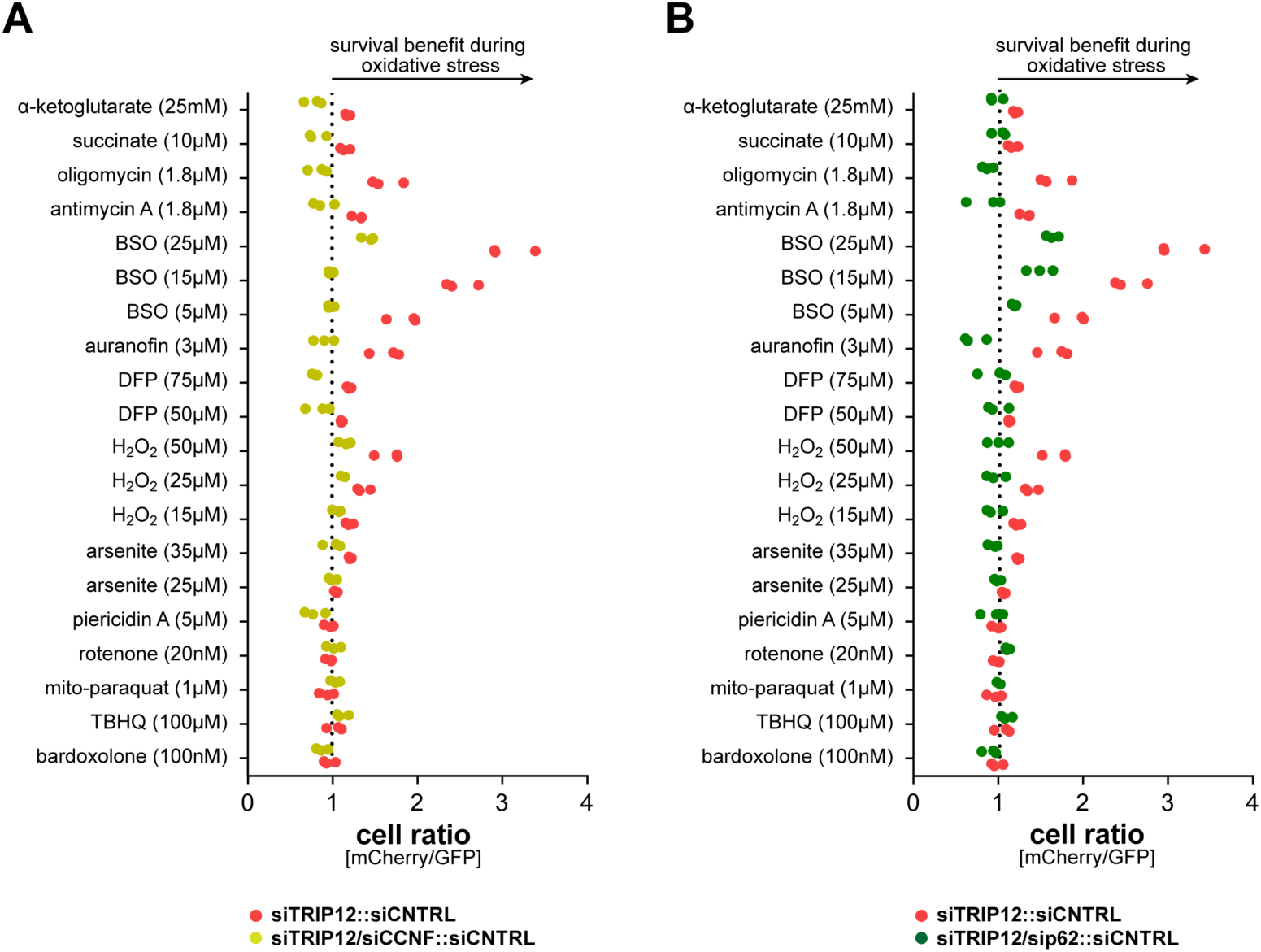
TRIP12 depletion promotes cell survival during stress. **A.** GFP-labeled control cells were mixed at a 1:1 ratio with mCherry-labeled cells depleted of TRIP12 (red dots). As indicated, CCNF was also depleted (yellow dots). Cells were exposed to increasing concentrations of oxidative stressors. After three days, the ratio of GFP- to mCherry-labeled cells was determined by flow cytometry. n=3 independent experiments. TRIP12-depleted cells are the same as shown in Figure 7B. **B.** GFP-labeled control cells were mixed at a 1:1 ratio with mCherry-labeled cells depleted of TRIP12 (red dots). As indicated, p62 was also depleted (green dots). Cells were exposed to increasing concentrations of oxidative stressors. After three days, the ratio of GFP- to mCherry-labeled cells was determined by flow cytometry. n=3 independent experiments. TRIP12-depleted cells are the same as shown in Figure 7B.

## STAR Methods

### EXPERIMENTAL MODEL AND STUDY PARTICIPTANT DETAILS

#### Cell Lines

All cell lines were purchased from ATCC. C2C12 myoblasts and HEK293T cells were grown in DMEM + glutamax (GIBCO, 51985091) with 10% fetal bovine serum (VWR, 97068-107). For C2C12 differentiation experiments, cells were grown to 70-90% confluence and media was changed into DMEM + glutamax (GIBCO, 51985091) with 2% fetal bovine serum (VWR, 97068-107). Media was changed daily for 3-4 days until cells differentiated. All cell lines regularly tested negative for mycoplasma.

### METHOD DETAILS

#### KEY RESOURCES TABLE

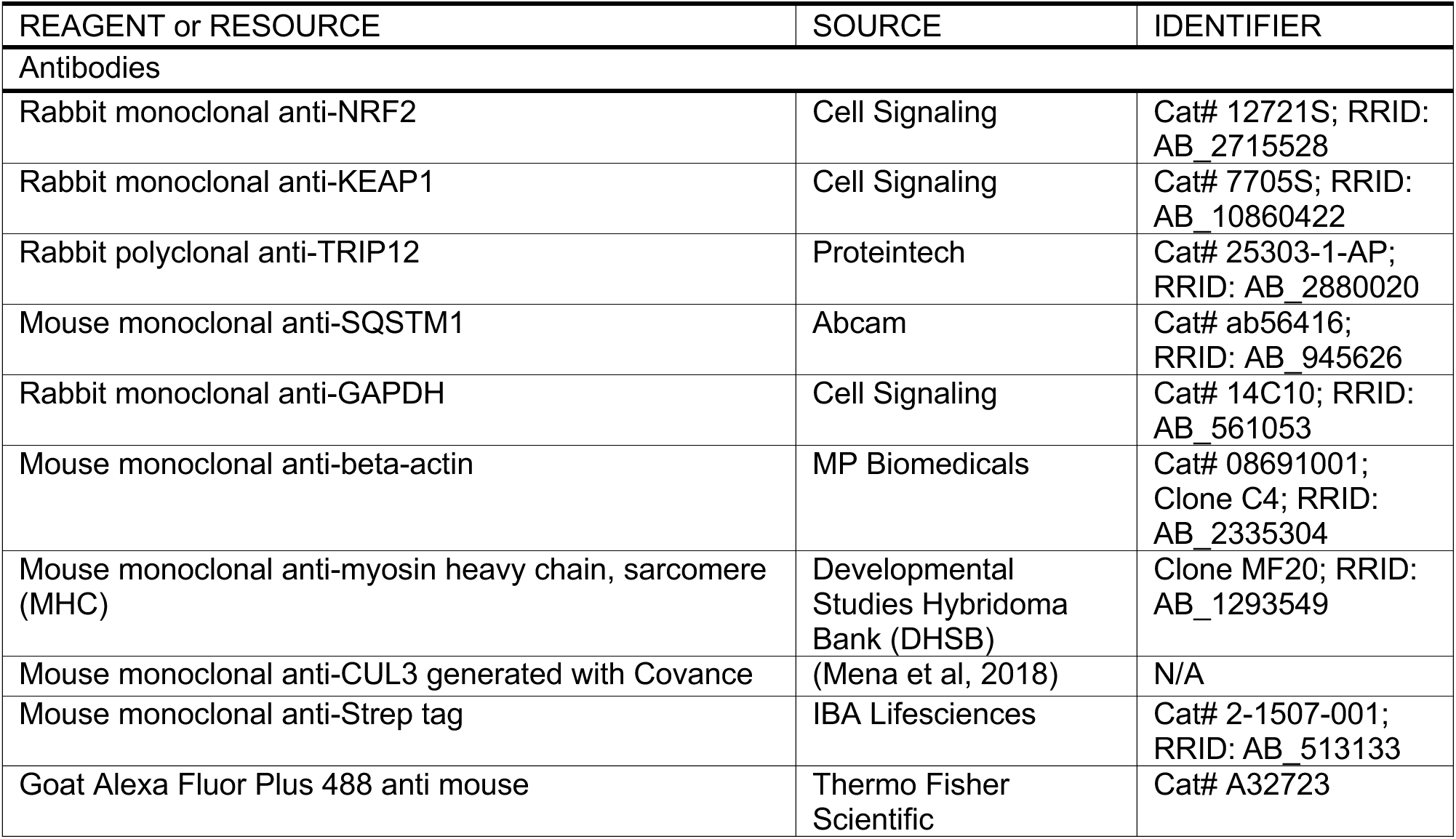
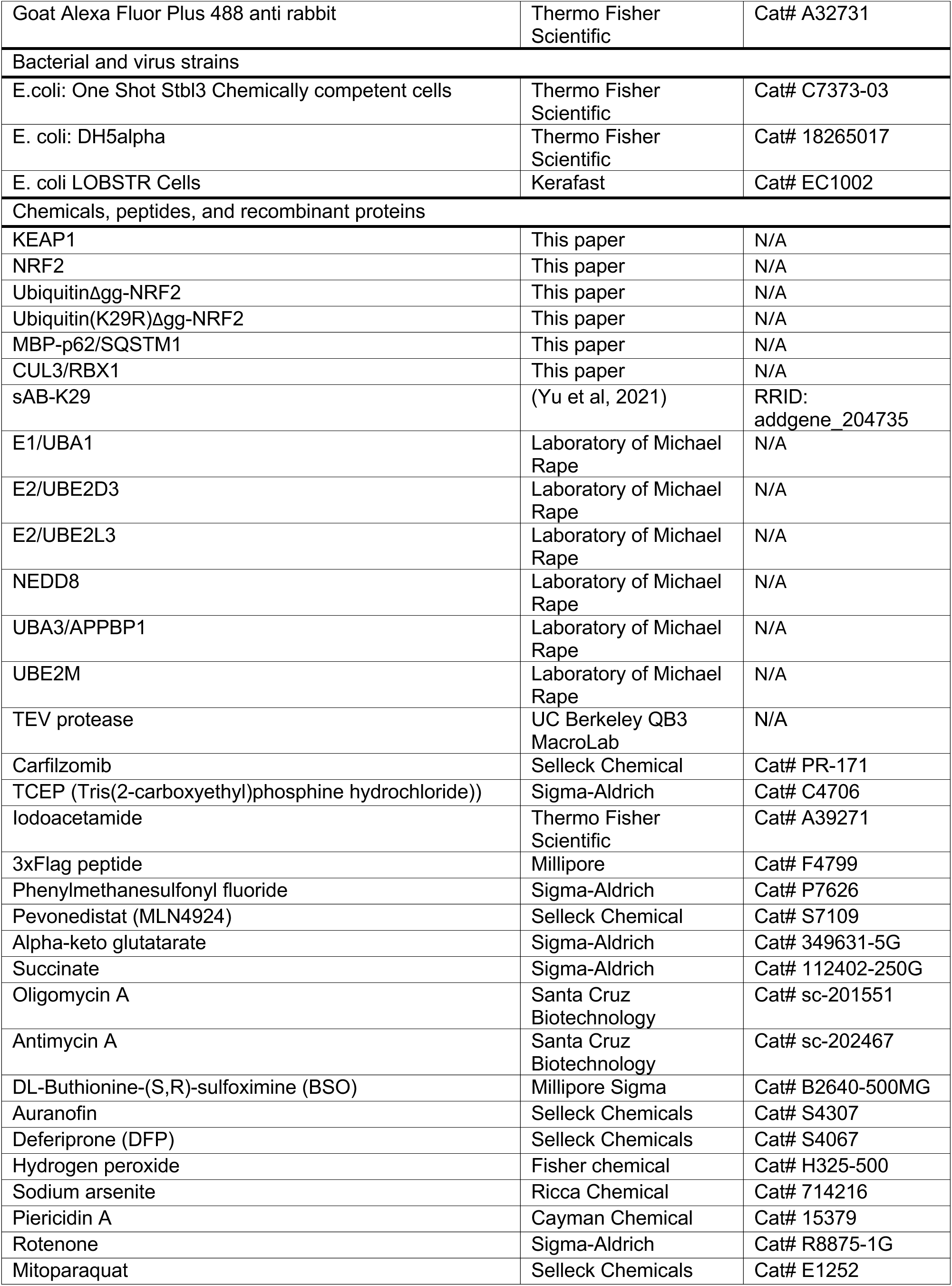
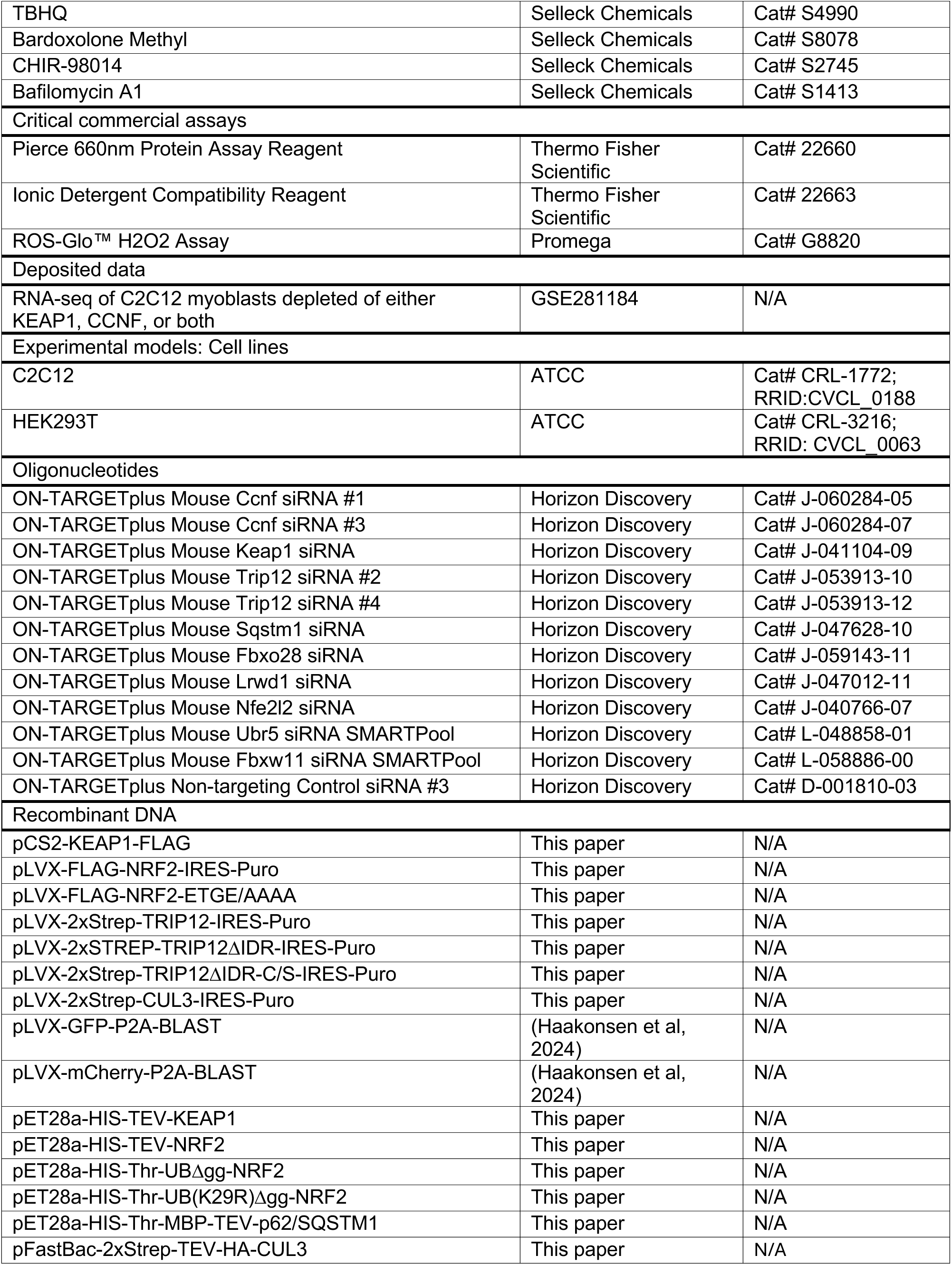
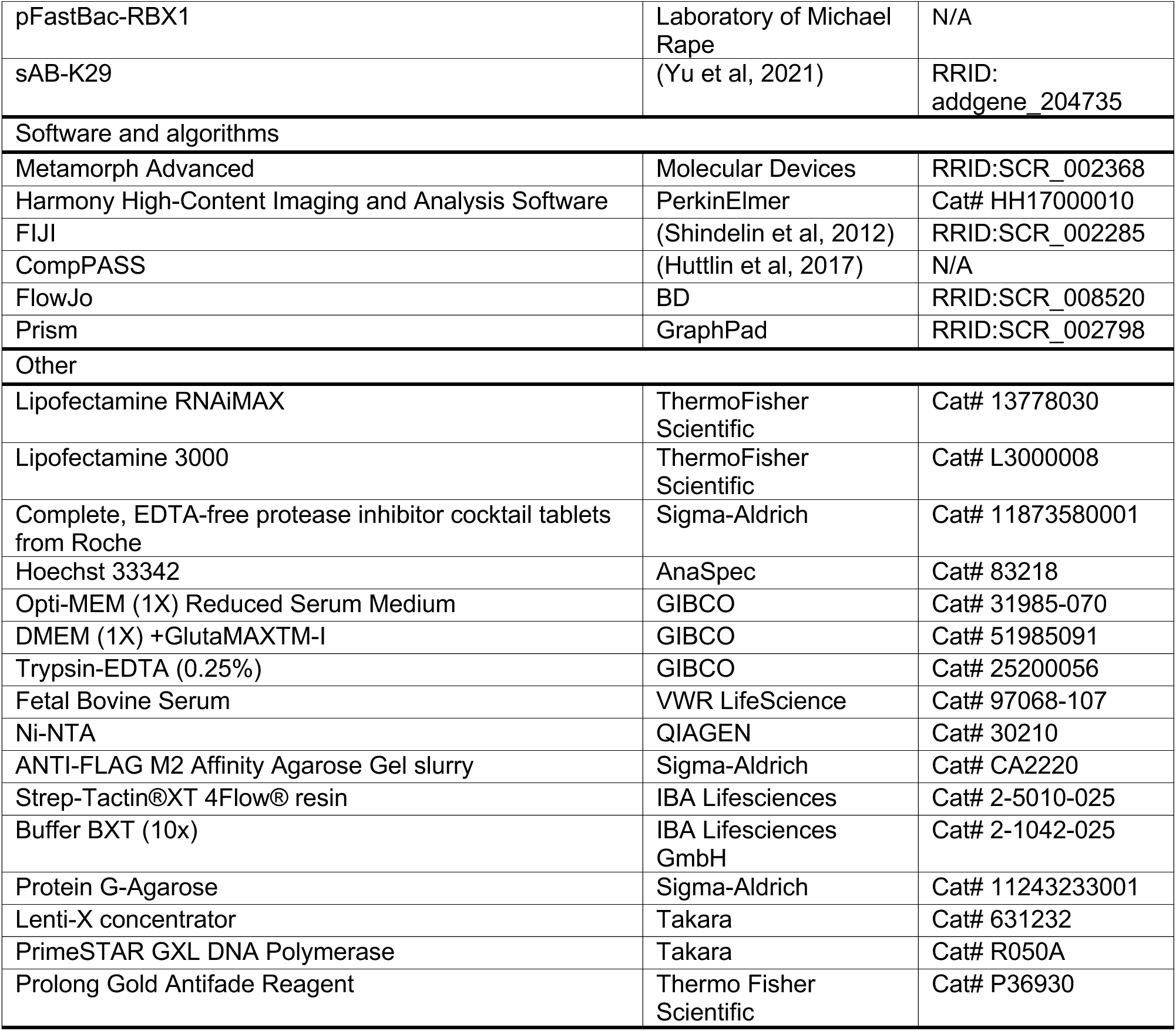

#### Cell Culture

C2C12 myoblasts and HEK293T cells were grown in DMEM + glutamax (GIBCO, 51985091) with 10% fetal bovine serum (VWR, 97068-107). C2C12 myoblasts were differentiated by changing media to DMEM + glutamax (GIBCO, 51985091) with 2% fetal bovine serum (VWR, 97068-107) daily for 3-4 days. siRNA transfections were performed with Lipofectamine RNAiMAX (Thermo Fisher Scientific, 13778030) according to manufacturer’s instructions. In C2C12 myoblasts, siRNAs were used at a final concentration of 30 nM. For experiments using western blotting or qPCR, cells were harvested 48h after siRNA transfection. All other transfections were done using Lipofectamine 3000 (Thermo Fisher Scientific, L3000008) according to manufacturer’s instructions.

#### Virus production and infection

Lentivirus for pLVX-2xStrep-CUL3-IRES-Puro, pLVX-GFP-P2A-Blast, pLVX-mCherry-P2A-blast, pLVX-3xFLAG-NRF2-IRES-Puro, and pLVX-3xFLAG-NRF2ΔETGE-IRES-Puro were made by co-transfection in HEK293T of viral plasmids with packaging plasmids using lipofectamine 3000 (Thermo Fisher Scientific, L3000008). Virus containing supernatants were collected 72h after transfection, spun down to remove any remaining cells, and concentrated in LentiX Concentrator (Takara, 631262) according to manufacturer’s instructions. After concentrating, virus was aliquoted and stored at −80°C for further use. To infect C2C12 myoblasts with virus, 50000 cells were seeded in a 12-well plate, mixed with virus and 6μg/ml polybrene (Sigma-Aldrich, TR-1003), and centrifuged for 45min at 1000g. Selection was started 24h later with either puromycin (1.5μg/ml, Sigma-Aldrich, P8833) or blasticidin (10μg/ml, Thermo Fisher Scientific, A1113903).

#### Immunoprecipitation and mass spectrometry

HEK293T cells were infected with pLVX-3xFLAG-NRF2-IRES-Puro or pLVX-3xFLAG-NRF2ΔETGE-IRES-Puro virus as described above. Cells were seeded in ten 15cm plates per condition and grown to 80-90% confluence. MLN4924 (500 nM Selleckchem, S7109) was added 16h before harvesting. Cells were lysed in 20 mM HEPES pH 7.5, 150 mM NaCl, 0.2% Nonidet P-40 (Sigma-Aldrich, E1014, 1x complete protease inhibitor cocktail (Roche, 11836170001), for 30 minutes at 4°C, clarified by centrifuging for 15min at 18000g at 4°C, and incubated with αFLAG M2 resin (Sigma-Aldrich, A2220) for 2h at 4°C. After incubation, resin was washed 3 times in lysis buffer and proteins were eluted with 0.5mg/ml 3xFLAG peptide (Sigma-Aldrich, F4799) buffered in 1x PBS, 0.1% triton X-100. Eluted proteins were precipitated overnight at 4°C by adding trichloroacetic acid to a concentration of 20%. Protein pellets were centrifuged and washed three times in acetone with 0.1M HCl, then dried at room temperature, and resuspended in 8M urea with 100mM Tris, pH8.5. Samples were reduced by adding 5mM TCEP (Sigma-Aldrich, C4706) for 20min, alkylated with 10mM iodoacetamide (Thermo Fisher Scientific, A39271) for 15min, diluted four-fold in 100mM Tris, pH8.5, and digested with 0.5mg/ml trypsin (Promega, v5111) with 1 mM CaCl_2_ overnight at 37°C while shaking. Samples were analyzed at the UC Berkeley Vincent J. Coates Proteomics and Mass Spectrometry Laboratory using multidimensional protein identification technology (MudPIT) and run on a LTQ XL linear ion trap mass spectrometer. High confidence interactors were identified using ComPASS to compare the identified peptides with numerous unrelated mass spectrometry samples from our laboratory ^95^. These high confidence interactors are presented in Figure 3D. Samples were normalized to the total spectral counts (TSC) of the NRF2 bait peptides. Complete proteomics data of all identified peptides is provided in Table S1.

#### Small-scale immunoprecipitation

Cells were grown in 1-3 15cm plates per condition and lysed as described. After clearing lysate by centrifugation, samples were normalized using Pierce 660 nm Protein Assay Reagent (Thermo Fisher Scientific, 22660). After normalization, 2.5% of the sample was saved as input and samples were incubated with either αFLAG M2 resin (Sigma-Aldrich, A2220) for KEAP1^FLAG^ pull-down or Strep-Tactin XT 4Flow resin (IBA, 2-5010-025) for ^Strep^CUL3 or Strep-NZF1 pull-downs at 4°C for 1h. Samples were washed three times in lysis buffer and eluted in twice in 120mM Tris, pH6.8, 4%SDS, M urea, 20% glycerol, 5% β-mercaptoethanol, bromophenol blue. SDS-PAGE and immunoblotting were performed using the indicated antibodies and images were captured using a ProteinSimple FluorChem M device.

#### Western blotting from whole cell lysates

Whole cell lysates were prepared by harvesting cells in lysis buffer as described above, but with the addition of 1μl Benzonase (EMD Millipore, 70746-4) per ml lysis buffer. Samples were incubated on ice for 15min and normalized as using Pierce 660 nm Protein Assay Reagent (Thermo Fisher Scientific, 22660). Samples were combined with 2x urea sample buffer and analyzed using SDS-PAGE and Western blotting using primary antibodies. Secondary antibodies conjugated to HRP were added after washing in 1x PBS with 0.1% Tween. Membranes were incubated for 2h, washed, and imaged by incubating with Immobilon Western Chemiluminescent Substrate (Millipore, WBKLS05000) according to manufacturer’s instructions and capturing images with a ProteinSimple FluorChem M device.

#### Recombinant DNA

Constructs were cloned into pCS2+, pLVX, pET28a, or pFastBac plasmid backbones using Gibson assembly using HIFI DNA Assembly master mix (NEB, E2621L) after amplification by PCR using PrimeSTAR GXL DNA Polymerase (Takara, R050A) of the gene of interest out of cDNA prepared from HEK293T cells using the Protoscript II First Strand cDNA Synthesis Kit (NEB, E6560S).

#### Antibodies

The following antibodies were used in this study: αNRF2 (Cell Signaling, 12721S), α KEAP1 (Cell Signaling, 7705S), αTRIP12 (Proteintech, 25303-1-AP), αSQSTM1 (Abcam, ab56416), αGAPDH (Cell Signaling, 14C10), α-β-actin (MP Biomedicals, 08691001, clone C4), αmyosin heavy chain, sarcomere (MHC) (mouse monoclonal; Developmental Studies Hybridoma Bank, clone MF20), αCUL3 (mouse monoclonal; generated with Covance ^96^), StrepMAB-Classic αStrep tag (IBA Lifesciences, 2-1507-001), Goat Alexa Fluor Plus 488 anti mouse (Thermo Fisher Scientific, A32723), and Goat Alexa Fluor Plus 488 anti rabbit (Thermo Fisher Scientific, A32731). sAB-K29 was used to identify K29-linked ubiquitin chains and was purified as previously reported ^72^. All antibody dilutions were determined experimentally.

#### siRNAs

The following siRNA reagents were used: ON-TARGETplus mouse Ccnf siRNA#1 (CCGCAGAGCUAUCGAAUCA), ON-TARGETplus mouse Ccnf siRNA#3 (CUACCGUGGUUGACUAUAA), ON-TARGETplus mouse Keap1 siRNA (GCGCCAAUGUUGACACGGA), ON-TARGETplus mouse Trip12 siRNA#2 (CGGCAGAGAGAUCCGGUUA), ON-TARGETplus mouse Trip12 siRNA#4 (CGCCUAGAUUGGAUAGAAA), ON-TARGETplus mouse Sqstm1 siRNA (GAACAGAUGGAGUCGGGAA), ON-TARGETplus mouse Fbxo28 siRNA (GCACAUUACAUACGGAUUU), ON-TARGETplus mouse Lrwd1 siRNA GACAAAGAGUCGAUGGGCU), ON-TARGETplus mouse Nfe2l2 siRNA (CAUGUUACGUGAUGAGGAU), ON-TARGETplus Non-targeting Control siRNA#3 (UGGUUUACAUGUUUUCUGA), ON-TARGETplus mouse Ubr5 siRNA SMARTPool (GUAUGAGAGUUUACGACAA, GUUCUUGACAUUCGGAUUU, ACUUGUAUUUCUCGACUUU, CGGUGGUACCUUAAAGAGA).

#### Recombinant Proteins

Human NRF2, KEAP1, and p62/SQSTM1 were cloned into pET28a with N-terminal 6xHIS tags (for p62/SQSTM1 6xHIS-MBP) followed by a TEV protease site to allow for tag removal. HIS-tagged proteins were purified from *E. coli* LOBSTR cells grown to OD 0.6 at 37°C and induced overnight at 16 C with 500μM IPTG. Cells were lysed in 50mM HEPES, pH7.5, 150mM NaCl, 1mM PMSF, 5mM β-mercaptoethanol, 10mg/ml lysozyme, 10mM imidazole) for 30min at 4°C, sonicated, and cleared by centrifuging at 20000g for 45min at 4°C. Cleared lysate was mixed with Ni-NTA slurry and incubated at 4°C for 2h. Ni-NTA resin was washed in 50mM HEPES, pH7.5, 150mM NaCl, 5mM β-mercaptoethanol, 20mM imidazole, three times for 15min each at 4°C. Proteins were eluted in 50mM HEPES, pH7.5, 150mM NaCl, 5mM β-mercaptoethanol, 250mM imidazole, and mixed with TEV protease overnight to remove tags. Proteins were dialyzed overnight into 50mM HEPES 7.5, 150 mM NaCl, 5 mM β-mercaptoethanol and re-incubated with Ni-NTA resin to remove tags and uncleaved proteins. Next, proteins were concentrated, analyzed for purity via Coomassie staining, and flash frozen for later use. Recombinant p62/SQSTM1 was never frozen and used for *in vitro* ubiquitylation assays immediately after purification and cleaving off the HIS-MBP tags. CUL3 and RBX1 were co-purified out of SF9 insect cells via strep pull down using a 2xStrep tag on CUL3. Strep pull downs are described above. RBX1 was untagged. After binding CUL3/RBX1 to strep resin and washing, CUL3/RBX1 was eluted using Buffer BXT Strep Elution Buffer (IBA Lifesciences, 2-1042-025), concentrated, analyzed for purity via Coomassie staining, and flash frozen for later use. The antibody recognizing K29 ubiquitin linkages (sAB-K29) was purified as described ^72^. E1 (UBA1), E2 (UBE2D3), E2 (UBE2L3), and neddylation machinery (UBA3/APPBP1, NEDD8, UBE2M) were purified as described ^97–99^. Ubiquitin and ubiquitin mutants were purchased from R&D Systems.

#### *In vitro* ubiquitylation

For *in vitro* ubiquitylations using TRIP12, pLVX-2xStrep-TRIP12 plasmids were transfected into three 15cm plates per condition and purified using Strep affinity-agarose as described above. For these reactions, all other components were added directly to strep resin that had bound TRIP12. All other *in vitro* ubiquitylations were performed in solution. Ubiquitylation assays were performed in a 10μl reaction volume with the following protein and buffer conditions: 1μM E1/UBA1, 1μM E2/UBE2D3, 1mg/ml ubiquitin (R&D Systems, U-100H), 10mM DTT, 1x energy mix (22.5mM creatine phosphate (Sigma-Aldrich, 10621714001-5G), 3mM ATP, 3mM MgCl_2_, 0.3mM EGTA, pH 7.5), 1× ubiquitylation assay buffer (25mM Tris, pH 7.5, 50mM NaCl, 10mM MgCl_2_), and 1μM substrate. PBS was used to fill to 10μl total volume. If CUL3^KEAP1^ were used in the reaction, 1μM NEDD8-modified CUL3/RBX1 and 1μM KEAP1 were added. CUL3/RBX1 were modified with NEDD8 by incubating in 25mM Tris, pH 7.5, 50mM NaCl, 10mM MgCl2, 1x energy mix (22.5mM creatine phosphate (Sigma-Aldrich, 10621714001-5G), 3mM ATP, 3mM MgCl_2_, 0.3mM EGTA, pH 7.5), 25μM Nedd8, 500μM DTT, 7μM CUL3 complexes, 500nM UBA3/APPBP1, 1μM UBE2M for 30min at 30°C. If p62 was used as a substrate, the MBP tag was cleaved overnight using TEV protease and TEV and uncleaved ^HIS-MBP^p62 were nickel subtracted away leaving only full-length p62/SQSTM1. After adding all components, *in vitro* ubiquitylations were incubated for 2h at 30°C with shaking, before the reaction was stopped by adding 2x urea sample buffer. Samples were analyzed by SDS-PAGE and Western. Ubiquitin linkage specificity was tested using commercially available recombinant human ubiquitin mutants (R&D Systems, UM-K6R, UM-K11R, UM-K27R, UM-K29R, UM-K33R, UM-K48R, UM-K63R, UM-NOK, UM-K60, UM-K110, UM-K270, UM-K290, UM-K330, UM-K480, and UM-K630).

For ^HIS^NRF2 purification after *in vitro* ubiquitylation, NRF2 was purified as described, but the HIS-tag was not removed after purification. All other enzymes used in this reaction had HIS or other purification tags removed to prevent isolation of unwanted proteins. After ubiquitylation, 5% of the reaction was saved as input, and the rest was diluted into 500μl 50mM HEPES, pH 7.5, 150mM NaCl, 5mM β-mercaptoethanol, and incubated with Ni-NTA resin for 1h at 4°C. After incubation, resin was washed three times in buffer, resuspended in 2x urea sample buffer and analyzed via Western blotting.

#### siRNA screens

C2C12 myoblasts were seeded into 96-well plates with 400 cells per well using a Thermo Fisher Scientific Multidrop Combi system. 24h later, cells were transfected with 30nM siRNA using an Agilent Velocity 11 Bravo Automated Liquid Handling Platform. 24h later, and each day for 4d, cells were differentiated by changing media once daily into fresh differentiation media. Then, cells were fixed in-well with 4% formaldehyde in 1x PBS for 20min, permeabilized with 0.1% triton in 1xPBS for 20min, blocked in 10% FBS in 1xPBS for 30min, and stained with an antibody recognizing myosin heavy chain and Hoescht33342. Myotubes were imaged on an Opera Phenix (PerkinElmer) using a 10x objective to capture 25 images per well. Images were analyzed using the PerkinElmer Harmony software to calculate the fusion index as previously described ^34^.

#### Myogenesis functional assays

C2C12 myoblasts were grown to 70-90% percent confluence in a 12-well plate, transfected with siRNAs, and media changed into differentiation media as described above. After 3-4d, cells were fixed in-well with 4% formaldehyde in 1x PBS for 20min, permeabilized with 0.1% triton in 1x PBS for 20min, blocked in 10% FBS in 1x PBS for 30min, and stained with primary antibody in 10% FBS and 1x PBS for 3h. After one wash in 1x PBS, secondary antibody and Hoescht33342 in 10% FBS and 1x PBS were added to cells for 1h. All steps were done at room temperature. Cells were imaged in-well using a Perkin Elmer Opera Phenix and images were analyzed using Harmony image analysis software.

#### Immunofluorescence microscopy

C2C12 cells were seeded at 5000 cells/ml on coverslips and transfected with siRNAs as described above. Cells were fixed with 4% formaldehyde in 1x PBS for 20min, permeabilized with 0.1% triton in 1x PBS for 20min, blocked in 10% FBS in 1xPBS for 30min, and stained with primary antibody in 10% FBS and 1x PBS for 3h. After one wash in 1x PBS, secondary antibody and Hoescht33342 in 10% FBS and 1xPBS were added to cells for 1h. All steps were done at room temperature. Coverslips were then mounted onto slides with Prolong Gold Antifade Reagent (Thermo Fisher Scientific, P36930) and imaged on an Olympus IX81 microscope equipped with a Yokogawa CSU-1X confocal scanner unit (CSUX1 Borealis Square Upgrade Module), ANDOR iXon3 camera (IXON DU-897-BV), and Andor Technology Laser Combiner System 500 series equipped with four laser lines. Images were analyzed in FIJI ^100^. For NRF2 nuclear localization, a mask was created using the Hoescht33342 stain to identify the nucleus and the average NRF2 signal intensity in the masked area was quantified. NRF2 nuclear localization experiments consist of at least 100 cells from two biological replicates. For p62 aggregation, the intensity of the p62 aggregates was multiplied by the area of the aggregates and then normalized to the number of cells in the frame. p62 aggregation experiments consist of three biological replicates.

#### RNA sequencing

C2C12 cells were transfected with siRNAs as described above and RNA of three biological replicates was extracted using a NucleosSpin RNA kit (Machery-Nagel, 740955). Library prep, sequencing, and analysis were performed by Novogene. Genes showing a greater than two-fold change compared to siCNTRL were kept for further analysis. Hierarchical clustering was performed and visualized by Morpheus (Morpheus, https://software.broadinstitute.org/morpheus). NRF2 targets ^101^ and E2F targets ^102^ were mapped onto the final heatmap after clustering.

#### DepMap

The Pearson correlations for all genes and either KEAP1 or CUL3 were downloaded from DepMap (https://depmap.org/portal) ^61^. Correlations were calculated using the DepMap Public 24Q2+Score, Chronos dataset. The complete list of correlations was compared to a list of E3 ubiquitin ligases ^103^to isolate ligases that may be related to KEAP1 and CUL3. Only positive correlations are shown.

#### ROS Measurements

Cellular levels of H_2_O_2_ were determined using the ROS-Glo™ H_2_O_2_ Assay (Promega, G8820) according to manufacturer’s instructions.

#### qPCR

C2C12 cells were transfected with siRNAs and RNA was extracted as described above. cDNA was synthesized using Protoscript II First Strand cDNA Synthesis Kit (NEB, E6560S). qPCR assays were done using 2xKAPA SYBR Fast qPCR master mix (Roche, KK4602) on a LightCycler 480 II Instrument (Roche). Expression fold changes were calculated using the ΔΔCt method.

#### Cell competition assays

C2C12 myoblasts were transduced to express either GFP or mCherry by infecting with pLVX-GFP-P2A-Blast or pLVX-mCherry-P2A-Blast viruses as described above. GFP-expressing cells were transfected with siCNTRL and mCherry-expressing cells were transfected with other siRNAs as described in Figures. 12-16h after siRNA transfection, GFP- and mCherry-expressing cells were counted and seeded with 50000 cells per well in a 12-well plate. 8h after seeding, drugs were added to each well. The ratio of GFP^+^/mCherry^+^ cells was determined 48h later using a BD LSRFortessa instrument, analyzed using FlowJo, and normalized to untreated siCNTRL sample. The ratio is calculated as ((siRNA_treatment_/siCONTROL_treatment_)/(siRNA_control_/siCONTROL_control_)) where siRNA_treatment_ is any non-siCNTRL sample.

### QUANTIFICATION AND STATISTICAL ANALYSIS

The quantifications presented in this study are shown as the mean ± standard deviation (SD). Myogenesis screens (**Figure 1A**; **Figure 2A**) are two technical replicates. All other myogenesis experiments are three biological replicates. RNA-seq analysis was done using three biological replicates. All qPCR is three technical replicates. NRF2 microscopy is at least 100 cells from two biological replicates and p62 microscopy is three biological replicates. All myogenesis experiments, immunofluorescence microscopy, and qPCR were analyzed for significance in GraphPad Prism by one-way ANOVA to compare all conditions at once (* p ≤ 0.05, ** p ≤ 0.01, *** p ≤ 0.001, **** p ≤ 0.0001).

